# Compartmentalization and synergy of osteoblasts drive bone formation in the regenerating fin

**DOI:** 10.1101/2023.06.04.543617

**Authors:** N Cudak, AC López-Delgado, F Rost, T Kurth, M Lesche, S Reinhardt, A Dahl, S Rulands, F Knopf

**Affiliations:** CRTD - Center for Regenerative Therapies TU Dresden, Germany; Center for Healthy Aging, Faculty of Medicine, TU Dresden, Germany; DRESDEN-concept Genome Center, DFG NGS Competence Center, c/o Center for Molecular and Cellular Bioengineering (CMCB), TU Dresden, Germany; Core Facility Electron Microscopy and Histology, Technology Platform, Center for Molecular and Cellular Bioengineering (CMCB), TU Dresden, Germany; Max Planck Institute for the Physics of Complex Systems, Dresden, Germany; Ludwig-Maximilians-Universität München, Arnold-Sommerfeld-Center for Theoretical Physics, München, Germany

**Keywords:** zebrafish, bone, single cell sequencing, lineage tracing, ablation, electron microscopy, osteoblast, blastema, osterix, matrix metallopeptidase 9, chondroitin sulfate, patterning, Fgf, Wnt

## Abstract

Zebrafish faithfully regenerate their fins after amputation which includes restoration of bone tissue and a component of cell plasticity. It is currently unclear how different cell populations of the regenerate divide labor to allow for efficient regenerate growth and proper patterning. Here, we studied lineage relationships of FACS-enriched epidermal, blastemal and bone forming fin regenerate cells by single cell (sc) RNA sequencing, lineage tracing, targeted osteoblast ablation and electron microscopy to show that the majority of osteoblasts in the outgrowing regenerate derive from *osterix*+ osteoblasts, while *mmp9*+ cells give rise to a limited cell number at the fin segment joints. A third population of distal blastema cells contributes to distal osteoblast progenitors, suggesting compartmentalization during appendage regeneration. Fin elongation and bone formation are carried out by distinct regenerate cell populations, and these variably depend on Fgf signaling. Ablation of *osterix*+ osteoblasts irreversibly impairs patterning of segment joints, and prevents bone matrix formation in the proximal regenerate. The resulting reduced regenerate length is partially compensated for by the distal regenerate which shows increased Wnt signaling activity. Surprisingly, ablation of joint cells does not abolish the formation of segment joints. Our study characterizes rare fin regenerate cell populations, indicates intricate osteoblast-blastema lineage relationships, inherent detection and compensation of impaired regeneration, and demonstrates zonation of the elongating regenerate. Furthermore, it sheds light on the variable dependence of bone formation on growth factor signaling.

## Introduction

Zebrafish rapidly regenerate complex tissues after loss, including their appendages, the fins. Due to fast bone restoration and transparency, the fin serves as a valuable tool to study bone regeneration ^1^. After amputation, a multi-layered wound epidermis (WE) forms, which is followed by blastema formation within 2 days post amputation (dpa) and subsequent fin outgrowth ^2^. The blastema, a mass of proliferative cells accumulating at the amputation plane subdivides into different zones: a distal most blastema (DMB) of several cell diameters in size and more proximal and lateral regions (proximal blastema), in which proliferation, patterning and osteoblast differentiation take place ^3^.

The descendance of osteoblasts in the regenerate from stump cell populations has been intensely studied. Mature *osterix*+/*osteocalcin*+ osteoblasts in the fin stump dedifferentiate, proliferate and migrate towards the forming blastema, where they contribute to restoration of bone matrices ^4–6^. Other, more progenitor cell-like stump cell populations add to the osteoblast cell pool in teleost fin regenerates as well, among them *mmp9*+ cells at the segment joints ^7, 8^. Whether and to what extend committed osteoblasts, osteoblast progenitors and non-osteoblast cells assembled *within* the early regenerate contribute to ongoing bone formation is unclear. In the regenerate, osteoblasts of different maturity reside in different locations. Osteoblast progenitors expressing *Runx2* localize to a region close to the DMB while differentiating *osterix*+ osteoblasts reside in more proximal positions. Fully mature osteoblasts expressing *osteocalcin* are found in proximity to the amputation plane at later stages of regeneration ^4, 9^. Thus, based on relative position within the regenerate and marker expression, a minimum of 3 osteoblast subtypes can be distinguished in the fin regenerate, although little is known about their respective expression profiles. The above *Runx2*/*osterix* hierarchy of osteoblasts in the distal regenerate appears to be supported by a pool of distal Runx2+ cells whose existence is maintained by Wnt/ß-catenin signaling and opposed by BMP signaling ^10^. While this awaits further investigation, delineation of lineage relationships between different osteoblast subtypes within the regenerate and their potential descendance from non-osteoblast sources is crucial to understand the basis of successful regeneration (with bone providing the necessary structural support to the regenerate) and to acknowledge the extent of cellular plasticity that is required for it.

In this study, we performed sc transcriptomics on fluorescence-activated cell sorted epidermal, blastema and osteoblast cell populations which may have been underrepresented in previous analyses ^11, 12^, due to the high abundance of outer epidermal and proximal mesenchymal populations in fin regenerates. We identified novel subpopulations and respective marker genes, and generated and tested hypotheses regarding lineage relationships of regenerate cell populations by performing trajectory inference and Cre-loxP genetic fate mapping. Tracing *osterix*+ osteoblasts and *mmp9*+ progenitor cells showed that both populations contribute to bone regeneration to different degrees, and that their ablation affects proliferation, patterning and matrix formation in different parts of the regenerate. Furthermore, trajectory analysis suggested that *shha*+ cells of the basal layer of the WE (BLWE) do not contribute to the osteoblast population, while Wnt-responsive *siam*+ cells in the distal blastema do so, as confirmed by label-retaining cell analysis. This progenitor cell population compensates for impaired regenerate elongation after impaired regeneration due to suppression of Fgf signaling or *osterix*+ cell ablation, a recovery process that is itself independent of Fgf-signaling but involves enhanced Wnt signaling. These findings, together with the identification of novel fin regenerate markers, advance our understanding of complex tissue regeneration in zebrafish.

## Results

### A robust regenerative response to repeated amputation

In order to enrich for DMB cells, osteoblasts and cells of the lateral BLWE of the growing 3 dpa fin regenerate, we FACS-isolated fluorescently labeled cells of quadruple transgenic reporter zebrafish, in which *siam*+, *Runx2*+, *osterix*+ and *shha*+ cells were labeled by either GFP or mCherry protein expression (**Fig. 1A, B**, **Suppl Fig. 1A**) and performed sc RNA sequencing. We repeated the procedure 4 times (Reg1-Reg4) using the same zebrafish in intervals of 4 weeks (**Fig. 1B**, **Suppl Fig. 1B**). Sc transcriptomic analysis across samples ^13^ revealed proliferating cells in distinct populations of the fin regenerate. We inferred the presence and UMAP location of the DMB by the near-absence of proliferating cells ^3^ and confirmed the presence of proliferating cells in other populations (*pcna, mki67,* **Fig. 1B, Suppl Fig. 1C**). Cluster composition revealed similar contributions of each amputation experiment (**Fig. 1B**) and showed that gene expression between the first (Reg1) vs the following (Reg2-4) samples was overall similar (**Suppl Fig. 1D)**. Significantly down-regulated transcripts were only detected for 5 genes with a fold change (FC) of less than 0.5, while no significantly upregulated genes with a FC beyond 2 were detected (**Suppl Fig. 1E**, **Suppl Table 1**). The overwhelming number of genes whose expression was similar between different samples (**Suppl Fig 2**) suggested that successive amputations do not influence gene expression levels in cells of the regenerate.

**Fig. 1.**
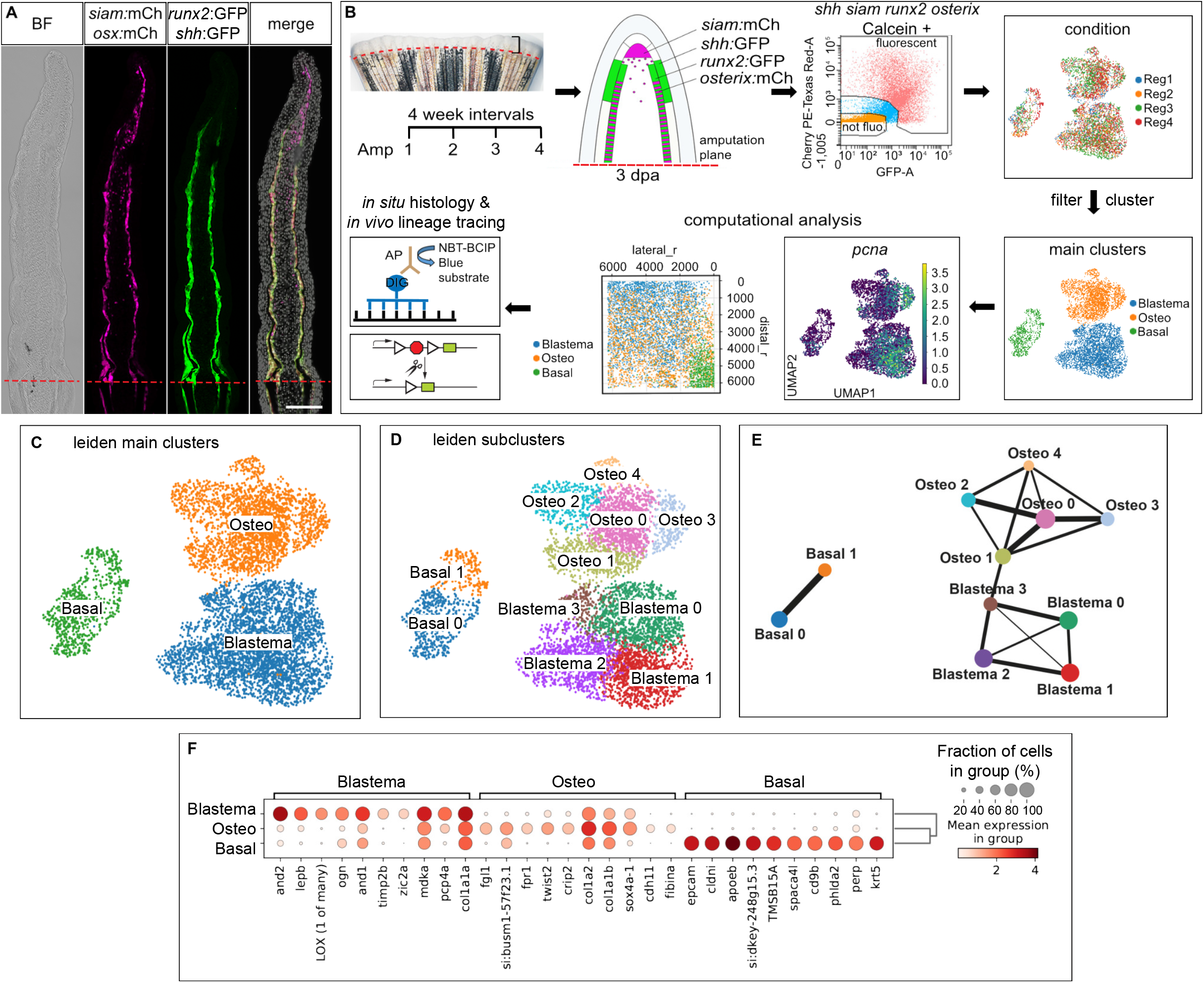
Experimental approach, the response to repeated amputation, cell clustering and trajectory analysis. (A) 3 dpa fin cryosection of quadruple transgenic reporter zebrafish. BF, brightfield, mCh, mCherry, *osx*, *osterix*. Scale bar 100 µm. (B) Study design. Reg, regeneration experiment, Osteo, Osteoblasts. Spatial reconstruction (pseudospace analysis): distal_r, pseudospace coordinate distal dimension, lateral_r, pseudo-space coordinate lateral dimension. AP, alkaline phosphatase coupled antibody, DIG, digoxygenin, NBT, nitro blue tetrazolium, BCIP, 5-bromo-4-chloro-3-indolyl-phosphate. (C) Main clusters identified in the analysis. Basal, BLWE; Blastema, blastema cells; Osteo, osteoblasts. (D) Identified subclusters. (E) PAGA analysis displaying connectivity (reflected by line thickness) between different clusters. (F) Marker gene expression in main clusters.

### Separation of epidermal from osteoblast and blastema cell clusters and their relative positions in the regenerate

In order to characterize the transcriptome of cell populations of interest, we used the combined dataset of Reg1-4 with 6668 cells for cluster analysis and identified 3 main clusters (**Fig. 1C**) encompassing 2, 4 and 5 subclusters, respectively (**Fig. 1D**). With the help of this dataset, we then identified marker genes for the respective subclusters and used these to localize them via RNA *in situ* hybrization. On a UMAP representation of the data, the main BLWE cluster (Basal) was clearly separated from the other two main clusters, which were connected and represented the osteoblast (Osteo) and blastema clusters, respectively (**Fig. 1D**). Trajectory inference using partition-based graph abstraction (PAGA) confirmed the Osteo-Blastema cluster connection (**Fig. 1E**) ^14^. We assessed the biological identity of the main clusters by inspecting marker genes (**Fig. 1F, Suppl Fig. 3A-C**, **Suppl Table 2**), which included *epcam, cldni, phlda2, fn1b, krt5* and *lef1* in the BLWE ^11, 15, 16^, *and1*/2, *lepb*, *her6*, *wnt5b* in the blastema ^17–20^ and *twist2*, *crip2* and *cdh11* in osteoblasts ^10, 11^ (**Fig. 1F, Suppl Table 2**). Gene set enrichment analysis uncovered overrepresentation of the GO terms peptide metabolic process, translation and peptide biosynthetic process in blastema cells (**Suppl Table 3**), ossification, skeletal system development and extracellular matrix organization in osteoblasts (**Suppl Table 4**) and cell adhesion, tight junction and cytoskeleton in cells of the BLWE (**Suppl Table 5**).

Next, we studied marker gene expression in the 2 BLWE, 4 blastema and 5 osteoblast subclusters (**Fig. 1D**). The subclusters showed specific, although not always exclusive, thus partly overlapping gene expression (**Suppl Fig. 3A-C, Suppl Tables 6-8**). Subcluster ‘Basal0’ encompassed more proximal proliferative BLWE cells expressing *col17a1b*, and *gstm.3* (**Suppl Fig. 3A, D**), while more distal ‘Basal1’ cells expressed *oclna*, *oclnb* and *fgf24* (**Suppl Fig. 3A, D**) ^21^. Blastema cells, categorized into Blastema0-3 subclusters expressed a variety of known and novel markers (**Fig. 1D, Suppl Fig. 3B, Fig. 2A, Suppl Fig. 4A**) ^20, 22–24^. Blastema0/1 cells represented proliferative cells positive for *postna*, *LOX* and *tnc* (**Fig. 1B, 2B, 2C, Suppl. 1C**, **3B**, **4B, Suppl Table 6**) corresponding to the highly proliferative proximal blastema ^3, 17^. Blastema2 cells were characterized by high expression of *aldh1a2* (**Suppl Fig. 4A**), *mmp13a* (**Suppl Fig. 4C**), *fgf10a* and *msx2b* (**Suppl Fig. 3C, Suppl Fig. 4A, Suppl Table 6**), therefore representing more distal blastema cells ^22–25^, and included non-proliferative *pcna-* and *mki67-* cells (**Fig. 1B, D, Suppl Fig. 1C**) expressing high levels of *lfng* ^26^, *mustn1a*, *mmp13a* ^27^, *bmp4* ^28^ and *tgfbi* (**Suppl Fig. 3B, Fig. 2D, Suppl Fig. 4C, Suppl Table 6**), corresponding to the DMB. Our analyses did not show exclusive markers for this non-proliferative region which suggests that the DMB is delimited on the basis of lacking proliferation rather than exclusive marker gene expression. Blastema3 cells, located in proximity to the main osteoblast cluster in the UMAP (**Fig. 1D**) showed high expression of *mfap5, pdgfrl* and *fhl1a* (**Suppl Fig. 3B, Fig. 2E, Suppl Table 6**).

**Fig. 2.**
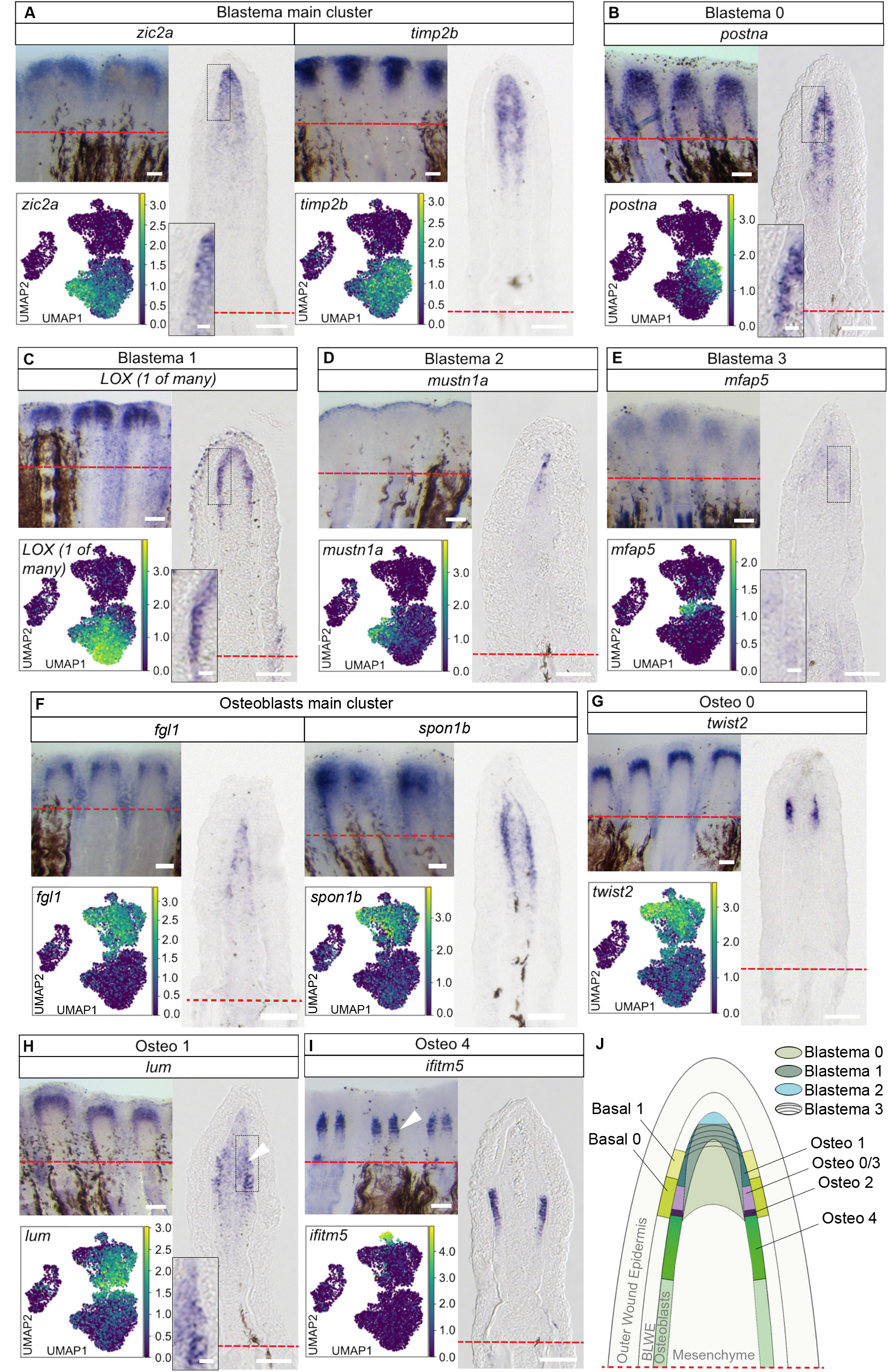
Phenotypic diversity and location of blastema cells and osteoblasts. (A) *zic2a* and *timp2b* expression. (B) *postna* in Blastema0. *postna* absence in osteoblasts and DMB. (C) *LOX (1 of many)* expression in Blastema1, touching the BLWE. (D) *mustn1a* expression. (E) *mfap5* expression in Blastema3 and some osteoblasts. (F) *fgl1* and *spon1b* expression. (G) *twist2* expression in Osteo0 cells. (H) Non-exclusive *lum* expression in Osteo1. (I) *ifitm5* expression. (J) Topology scheme of the 3 dpa regenerate, with vague distinction between Blastema0, 1 and 3. (A)-(I) UMAP, whole mount RNA ISH (WMISH) and cryosection views. Scale bars whole mounts 100 µm, cryosections 50 µm, insets 10 µm.

Bone forming cells expressed *fgl1*, *twist2* and *spon1b* (**Fig. 1F, 2F, 2G, Suppl Table 2**), and comprised 5 subclusters (**Fig. 1D, Suppl Fig. 3C, Suppl Fig. 4D-I**), characterized by often overlapping gene expression (**Fig. 1F, Suppl Fig. 3C**). Osteo0, a central cluster with a high number of cells, was positive for *twist2, twist3* and *tnc* expression (**Suppl Fig. 3C, Suppl Fig. 4B, Fig. 2G**). Osteo1, the cluster found close to Blastema3 cells in the UMAP (**Fig. 1D**) showed *lum* and *sgk1* expression (**Fig. 2H, Suppl Fig. 4E**). The expression of *ednrab, evx1, cx43*, *mmp9*, *hoxa13a* and *pthlha* ^15, 29–31^ clearly defined the Osteo2 population as joint cells (**Suppl Fig. 3C, Suppl Fig. 4F**, **Suppl Table 7**), while Osteo3 cells denoted proliferative osteoblasts positive for *stmn1a* (**Suppl Fig. 4G,** compare UMAP with *pcna* and *mki67* UMAPs in **Fig. 1B, Suppl Fig. 1C**). Osteo4 expression was characterized by high abundance of *osterix*, *bgna*, *spp1*, *col10a1a* and *entpd5a* transcripts (**Suppl Fig. 4H, Suppl Table 7**), which are found in differentiating zebrafish fin regenerate osteoblasts ^4, 30, 32^. High *mCherry* transcription (**Suppl Table 7**) relative to the other 4 subclusters and the expression patterns of the markers *ifitm5, panx3, sgm2* and *fgfbp2* (**Fig. 2I, Suppl Fig. 4I**) confirmed that Osteo4 represented differentiating osteoblasts. We delineated the relative subcluster positions by comparing mRNA *in situ* hybridization (ISH) patterns, leading to a distal regenerate topology of cells that we have enriched for (**Fig. 2J**). The most proximal population of analyzed osteoblasts are differentiated *osterix:mCherry*+ osteoblasts labeled by *panx3, sgms2, fgfbp2a* and *ifitm5* (**Fig. 2I, J, Suppl Fig. 4I**). Distal to this Osteo4 zone of differentiated osteoblasts (**Fig. 2J, Suppl Fig. 5A**), occasionally within the Osteo4 zone (arrowheads in **Fig. 2I, Suppl Fig. 4I**) Osteo2 joint cells are found (**Suppl Fig. 4F, Suppl Fig. 5B**). Distal to the Osteo4-Osteo2 region, highly proliferative *stmn1a*+ /*kpna2*+ Osteo3 cells are found (**Fig. 2J, Suppl Fig. 4G, Suppl Fig. 5C**), together with *twist2* and *tnc* double+ Osteo0 cells (**Suppl Fig. 5D**). *lum*+, *sgk1*+, (**Fig. 2H, Suppl Fig. 4E**), *abi3bpb*+*, mxra8b*+ (**Suppl Fig. 4D**) Osteo1 cells represent the distalmost osteoblast population within the regenerate.

Blastema2 cells encompass DMB cells. Underneath, proliferative peripheral (touching the BLWE) Blastema1 cells enriched for *LOX (1 of many)* (**Fig. 2C**) and more central (devoid of contact with BLWE) *postna*+ Blastema0 cells (**Fig. 2B**) can be found. *mfap5*+ Blastema3 cells (**Fig. 2E**) reside in a position directly adjacent (distal) to Osteo1 osteoblasts (**Fig. 2J**). Both populations share expression of *abi3bpb* and *mxra8b* (**Suppl Fig. 4D**) and might thus represent a transition zone of blastema cells and osteoblasts. The diversity of osteoblast, blastemal and BLWE clusters in our dataset illustrates the advantage of FACS-enriching for rare cell populations of interest.

### Lineage tracing of osterix+ and mmp9+ osteoblasts suggests variable contribution to bone formation and the presence of different regenerate domains

Next, we made use of CreERT2-loxP-mediated lineage tracing (**Fig. 3A**) to determine the contribution of differentiated *osterix*+ osteoblasts (including Osteo4 osteoblasts, **Fig. 3B-D**) and *mmp9*+ osteoblasts (including Osteo2 and likely Osteo1 osteoblasts, **Fig. 3E-G**) as the fate and extent of progeny formation of these osteoblast populations *within* the elongating regenerate were unexplored. We activated CreERT2 in *osterix*:CreERT2-p2a-mCherry x *hsp70l*:R2nlsGFP ^33^, *mmp9*:CreERT2 x *hsp70l*:R2nlsGFP, and *osterix*:CreERT2-p2a-mCherry x *mmp9*:CreERT2 x *hsp70l*:R2nlsGFP zebrafish fin regenerates, respectively, by injection of 4-hydroxytamoxifen (4-OHT) at 2.5 dpa, when CreERT2 is expressed in osteoblasts/ osteoblast progenitors of the regenerate ^4, 7^ (**Fig. 3A**). *osterix*+ recombined, nuclear GFP (nGFP)+ cells in *osterix*:CreERT2-p2a-mCherry x *hsp70l*:R2nlsGFP zebrafish were dispersed in the 3 dpa regenerate, their location spanning the region of Osteo4 cells (**Fig. 3B**) and more proximal osteoblasts, in which CreERT2 and mCherry protein is produced in the transgenic Cre-driver zebrafish (**Suppl Fig. 6**). At 4 and 5 dpa the regenerate had considerably and progressively enlarged, as did the nGFP+ domain (62,45 ± 5 % of the regenerate at 3 dpa, 79,46 ± 7,8 % at 4 dpa and 84,18 ± 3,5 % at 5 dpa, **Fig. 3C**). At this time, the distalmost nGFP labeled cells in the regenerate were detected ∼250 µm apart from the regenerate tip (compared to ∼350 µm at 3 dpa), with the density of nGFP labeled regenerate cells remaining constantly high (**Fig. 3D**). This suggests profound contribution of *osterix*+ osteoblasts and their progeny to bone forming cells in proximal regions of the regenerate, sparing the most distal 250 µm of the regenerate. In *mmp9*:CreERT2 x *hsp70l*:R2nlsGFP zebrafish, the progeny of recombined *mmp9*+ cells labeled by nGFP was found close to the joints, but also represented scattered cells within the fin rays and sometimes interrays (**Fig. 3E**). While recombined cells at the joints neither drastically changed their position (arrowheads in **Fig. 3E**) nor number (average clone size 24 ± 10 cells, **Fig. 3F**) until 5 dpa, distal scattered cell clones amplified in size (**Fig. 3E**), varying considerably in number (average clone size 54 ± 42 cells, **Fig. 3G**). This indicates a limited contribution of *mmp9*+ cell progeny at the prospective joints to the osteoblast regenerate populations, while distal *mmp9*+ cells may contribute to considerably more osteoblasts, but also non-osteoblast tissue in the regenerate. In *osterix*:CreERT2-p2a-mCherry x *mmp9*:CreERT2 x *hsp70l*:R2nlsGFP zebrafish, in which both *osterix*+ and *mmp9*+ cells had been recombined, more cells were labeled by nGFP than in the individual Cre-driver line experiments. This was evident by a larger domain of nGFP+ cells in individual fin rays of zebrafish in which both Cre-drivers were present (**Fig. 3H**) and a concomitant shorter distance of labeled cells to the regenerate tip (brackets in **Fig. 3I**). These data indicate that *osterix*+ cells including Osteo4 cells and *mmp9*+ cells including Osteo2 and Osteo1 cells contribute to different sets of fin regenerate osteoblasts in different compartments and at different quantities.

**Fig. 3.**
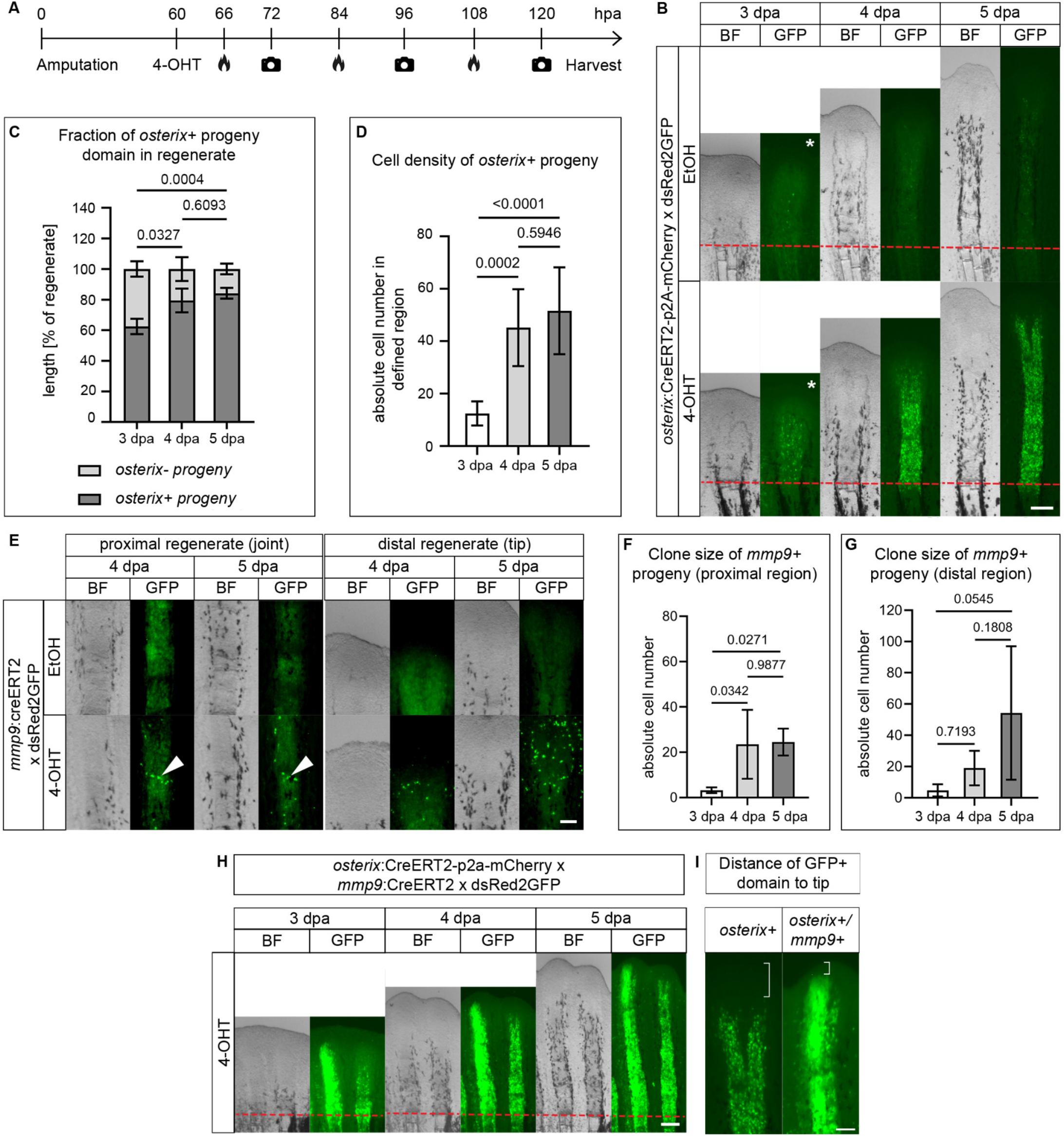
Lineage tracing of *osterix*+ and *mmp9*+ osteoblasts. (A) Experimental design. 4-OHT, 4-hydroxytamoxifen. Flame icon, heat induction. Camera icon, imaging. (B) Lineage tracing of *osterix*+ cells. Asterisk, brightness and contrast increased to reveal GFP expression. Scale bar 200 µm. (C) Fraction of *osterix*+ and *osterix*-progeny at 3, 4 and 5 dpa. Kruskal-Wallis. (D) Density of *osterix*+ progeny in the 3, 4 and 5 dpa regenerate. Defined region = 0.028 mm^2^. One-way Anova (Tukey). (E) Lineage tracing of *mmp9*+ cells [joint cells in proximal regenerate (arrowhead), left panel, and cells in distal regenerate, right panel]. Scale bar 100 µm. (F) Clone size of *mmp9*+ progeny in the proximal region of the regenerate. One-way Anova (Tukey). (G) Variable clone size of *mmp9*+ progeny in the distal region of the regenerate. One-way Anova (Tukey). (H) Lineage tracing of *osterix*+ and *mmp9*+ cells. Scale bar 200 µm. (I) Magnified view of *osterix*+ progeny shown in (B) and *osterix*+/*mmp9*+ progeny shown in (H). Brackets, distance to the tip of the regenerate. Scale bar 100 µm. (B), (E), (H) EtOH, ethanol vehicle control. Red dashed line, amputation plane. BF, brightfield.

### Mixing of blastema and osteoblast cell populations in the distal regenerate suggests contribution of distal blastema cells to bone formation

The fact that the progeny of *osterix*+ cells spared a region at the regenerate tip indicated that other cells may contribute to distal osteoblast populations. In order to explore this possibility, we performed transmission electron microscopy (TEM) of 3 dpa regenerating fins and compared cell phenotypes of WE-underlying blastema and osteoblast cells in different proximo-distal positions of the regenerate (**Fig. 4A, B, Suppl Fig. 7**). While DMB cells were small with few endoplasmatic reticulum (ER) and mitochondria and produced a thin ECM layer, putative pre-osteoblasts and osteoblasts produced more ECM and showed a more secretory phenotype with more dilated ER Golgi complexes and mitochondria especially close to the amputation plane (**Fig. 4B, Suppl Fig. 7**). We did not discern any abrupt phenotype changes between cells but rather a gradual change in morphology and tested for cells transitioning between the distal and proximal blastema. A prime candidate for a transitioning population was the *runx2*:GFP+ cell population (**Fig. 1A**) ^9, 10^, in addition to distal *mmp9*+ cells. Of note, *runx2a* transcripts were detected particularly in proliferating osteoblasts (Osteo0, Osteo3) (**Fig. 4C**), while its expression was low in Osteo4 cells. We tested whether distal *runx2*:GFP+ osteoblasts are generated by blastema cells by using transgenic *siam*:mCherry x *runx2*:GFP zebrafish, in which DMB cells and premature osteoblasts are labeled by *mCherry* and *gfp* transcripts and proteins, respectively (**Fig. 4D-F**). Live imaging, FACS and combined RNA ISH and immunohistochemistry (ISH-IHC) were used to follow the respective cell progeny. Double RNA ISH showed that *gfp* and *mCherry* reporter transcript domains were sometimes clearly separated but often continuous with a slight overlap (arrowhead in **Fig. 4E**), as previously described ^25^. ISH-IHC revealed that *siam*:mCherry protein distribution extended much more proximally than *mCherry* transcripts, and that there was only a minimum overlap between *mCherry* mRNA and GFP protein (arrowheads in **Suppl Fig. 8A**). We hypothesized that some of these *siam*:mCherry protein+ cells had lost *siam* promotor activity and accordingly *mCherry* transcription, exited the distal Wnt-active domain and relocated proximally to contribute to the osteoblast cell pool. In order to test this, we performed FACS analysis in *runx2*:GFP x *siam*:mCherry fin regenerates and detected a considerable number of cells with simultaneous red and green fluorescence, i.e. mCherry and GFP protein overlap (**Fig. 4D**). Confocal imaging of 3 dpa regenerates of the same zebrafish line revealed that GFP and mCherry double+ cells were detectable beyond 150 µm proximal to the distalmost DMB mCherry+ cells (protein level, **Fig. 4F, G)**. We never detected *mCherry* transcripts in such proximal locations, and concluded that *siam*:mCherry+ blastema cells (potentially reflecting *and1*+, *and2*+ actinotrichia-forming cells, **Suppl Fig. 8B**) ^19^ contribute to the pool of premature osteoblasts in the regenerate. We also detected some cells with low GFP protein levels outside of the domain of *gfp* transcription in the tip region (0-150µm, **Suppl Fig. 8C**), suggesting that *runx2*:GFP+ pre-osteoblasts contribute to or mix with the distal blastema cell population. We next tested whether there was any overlap between *siam*:mCherry transcripts and *gfp* transcripts in *osterix*:GFP x *siam*:mCherry zebrafish using the same approach, with *osterix*:GFP+ cells being more proximally located than *runx2*:GFP+ cells. *gfp* and *mCherry* transcript domains were clearly separated (**Fig. 4H**). Nevertheless, GFP and mCherry double+ cells were detected proximal to *siam*:mCherry transcript expression (**Fig. 4G, I, Suppl Fig. 8A**). These observations support the hypothesis that distal Wnt-active cells contribute to the osteoblast cell pool in the distal regenerate.

**Fig. 4.**
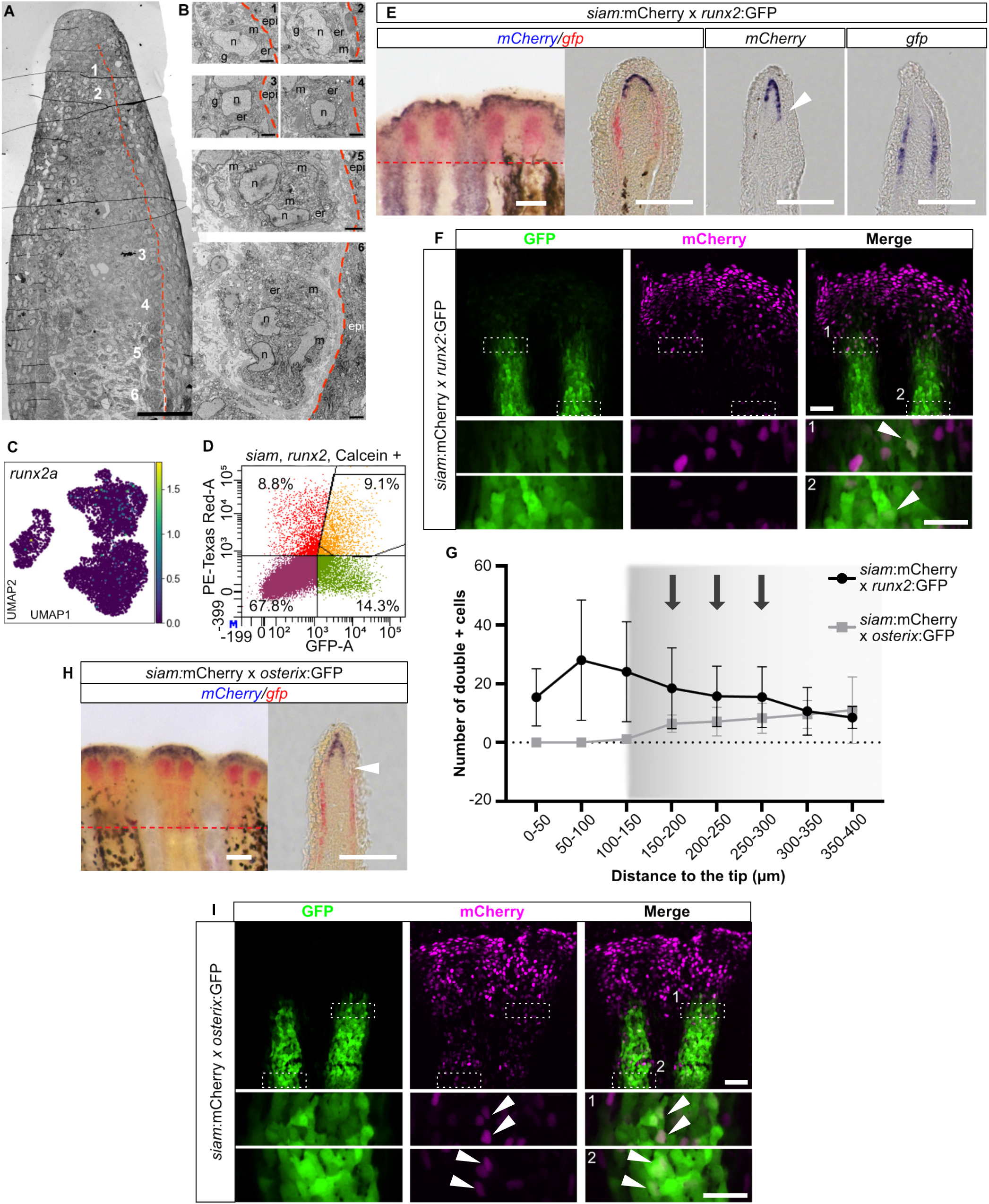
Mixing of blastema and osteoblast cells. (A) Overview of TEM of regenerate with positions of cells of interest indicated (1-6). Red dashed line, subepithelial basal lamina. (B) Cells underlying the BLWE with gradual increase of endoplasmic reticulum (er) from distal (1,2) via intermediate (3,4) to proximal positions. More dilated er in proximal regions with more Golgi complexes (g) and mitochondria (m) suggesting massive protein synthesis. epi, epidermis. n, nucleus. Scale bar (A) 50 µm, (B) 2 µm. (C) UMAP of *runx2a*. (D) FACS gates to identify *siam*:mCherry+, *osterix*:GFP+ and mCherry/GFP double+ cells at 3 dpa with respective percentages of unlabeled and labeled cell populations. (E) Single and double ISH of *mCherry* and *gfp* transcripts in *siam*:mCherry x *runx2*:GFP transgenic zebrafish. Arrowhead, proximal *mCherry* expression. Red dashed line, amputation plane. Scale bars 100 µm. (F) Detection of mCherry/GFP protein double+ cells (arrowheads in boxed areas) in 3 dpa fin regenerates of *siam*:mCherry x *runx2*:GFP transgenic zebrafish. Scale bar overview 50 µm. Scale bar inset 25 µm. (G) Quantification of mCherry, GFP double+ cells in *siam*:mCherry x *runx2*:GFP and *siam*:mCherry x *osterix*:GFP transgenic zebrafish [experiments shown in (F), (I)]. Shadowed interval with arrows, regions outside of transcript detection. (H) Double ISH of *mCherry* and *gfp* transcripts in *siam*:mCherry x *osterix*:GFP transgenic zebrafish. Arrowhead, transcript negative region. Red dashed line, amputation plane. Scale bars 100 µm. (I) Detection of mCherry/GFP protein double+ cells in 3 dpa *siam*:mCherry x *osterix*:GFP fin regenerates (arrowheads in boxed areas). Scale bar overview 50 µm. Scale bar inset 25 µm.

### Fgf signaling controls osteoblast and distal blastema compartment sizes

Next, we tested whether signaling pathway alterations would affect the size and function of these different regenerate domains. We turned to Fgf signaling, which is known to impact skeletal growth in various species ^34^ and which is active in regenerating fins ^24, 35, 36^. Suppression of Fgf signaling by SU5402 in *siam*:mCherry x *osterix*:GFP zebrafish from 3 to 5 dpa led to an overall reduced regenerate length (**Fig. 5A**, arrowhead in **Fig. 5B**). This reduction was attributable to a shortened *osterix*+ domain, while neither the size and Wnt reporter activity of the *osterix*-, *siam*+ tip region nor the activity of the *osterix*:GFP+ cells were affected (**Fig. 5A-C, Suppl Fig. 9**). We let fins recover from the inhibitor treatment to test whether Fgf suppressed regenerates would catch up in growth. Indeed, fins recovered regenerate length during a two-day recovery period in which both the size and the reporter activity of the distal Wnt active domain increased (**Fig. 5D-F**). Furthermore, *osterix*:GFP signal intensity was higher in regenerates recovering from Fgf suppression, suggesting enhanced osteoblast maturation during catch up (**Fig. 5F**). These results indicate that the Wnt+ domain encompassing DMB cells and pre-osteoblasts compensate for impaired regenerate growth after Fgf signaling suppression which cannot erase the presence of positional identity in the regenerate.

**Fig. 5.**
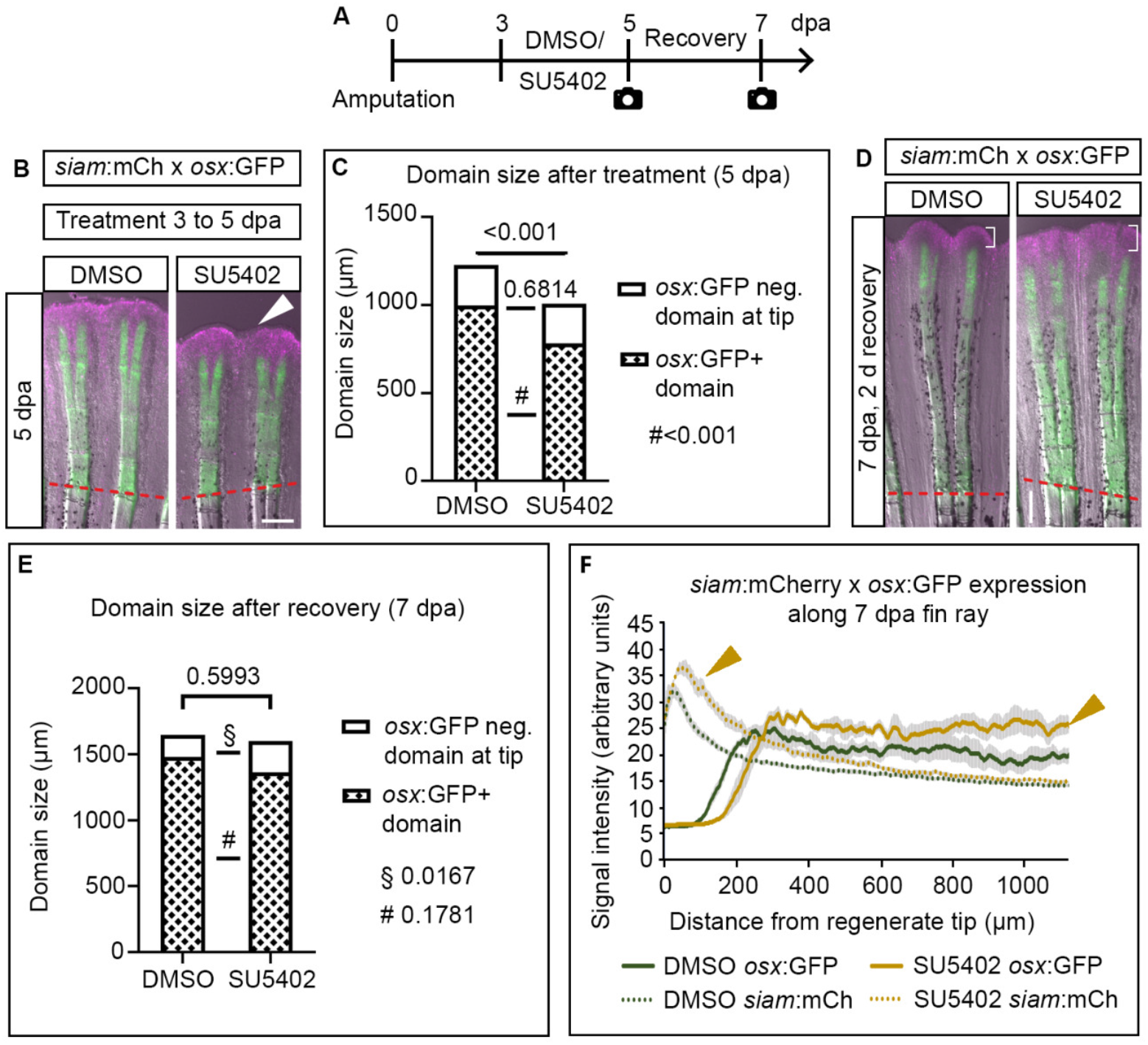
Effects of SU5402 on different regenerate domains. (A) Treatment regimen. (B) 5 dpa fin regenerates of *siam*:mCherry x *osterix*:GFP transgenic zebrafish treated with SU5402 or DMSO from 3 to 5 dpa. (C) Quantification of experiment shown in (B). Welch’s t-tests. (D) 7 dpa fin regenerates of *siam*:mCherry x *osterix*:GFP transgenic zebrafish treated with SU5402 or DMSO from 3 to 5 dpa and 2 day recovery (5 to 7 dpa). Brackets, *siam*+ tip region. (E) Quantification of experiment shown in (D). Welch’s t-tests. (F) Fluorescence signal intensity of transgenic reporters along fin regenerate. Mean ± SEM. Arrowheads, increased signal intensity. Scale bars 200 µm.

### Osteoblast ablation impairs regeneration domain-specifically and regenerates can partially recover

We further investigated the contribution of different osteoblast populations to bone regeneration by ablating either *osterix*+ cells (proximal regenerate), *mmp9*+ cells (distal regenerate), or both cell populations simultaneously. Transgenic zebrafish expressing nitroreductase (NTR) in the respective cells ^7, 37, 38^ were treated with the substrate nifupirinol (NFP) ^39^, resulting in toxic product formation in osteoblasts. *osterix*:NTR/*mmp9*:NTR double ablation from 3 to 5 dpa reduced regenerate length significantly when compared to individual *osterix*+ and *mmp9*+ cell ablation (**Fig. 6A, B**). Individual *osterix*:NTR+ cell ablation had a stronger anti-regenerative effect than *mmp9*:NTR+ cell ablation. This confirmed the importance of *osterix*+ cells for bone regeneration, and corroborated the finding that *osterix*+ and *mmp9*+ cells represent distinct fin regenerate cell populations.

**Fig. 6.**
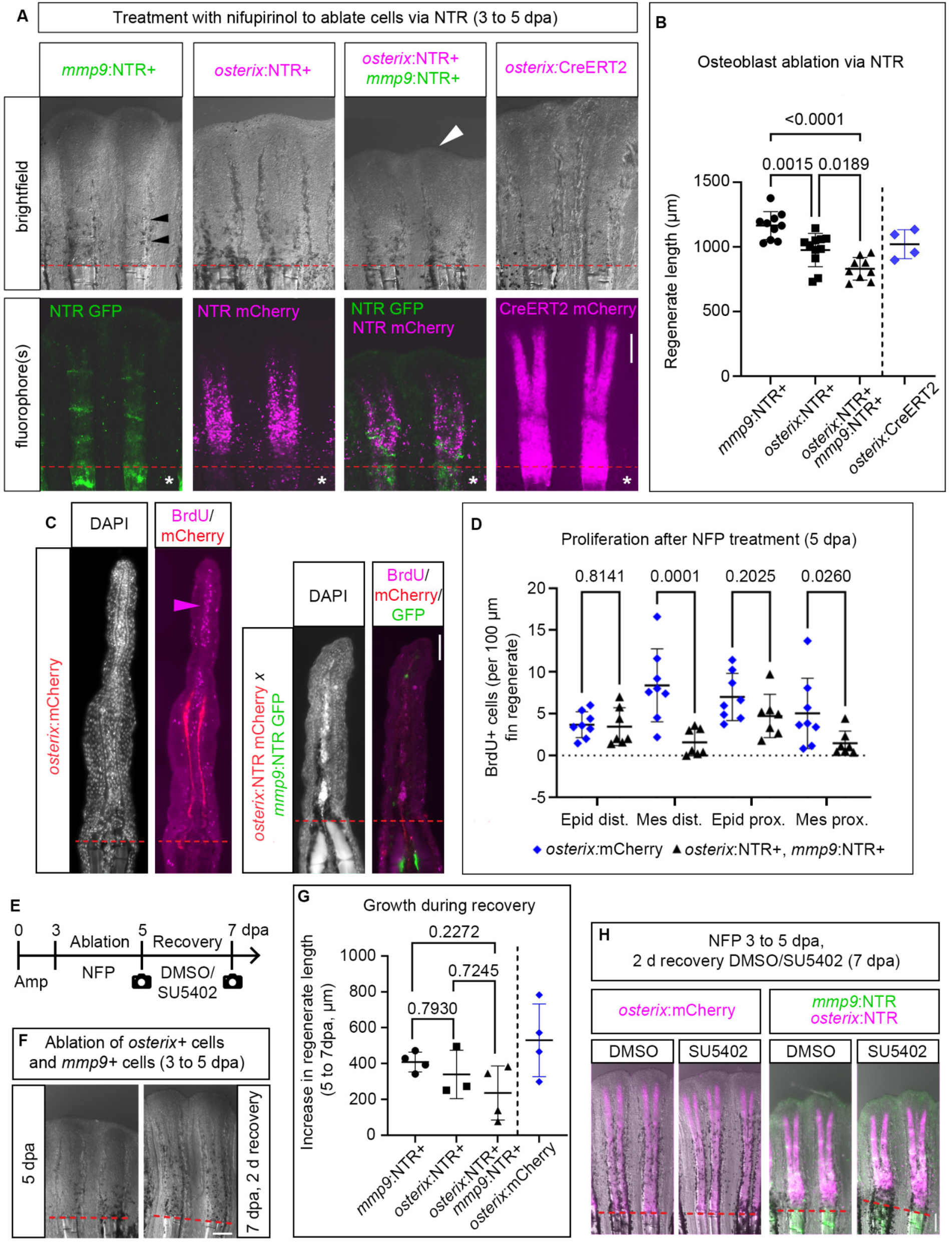
Effects of *mmp9*+ and *osterix*+ cell ablation on fin regeneration. (A) 5 dpa fin regenerates of 3 to 5 dpa NFP treated fin regenerates of *mmp9*:NTR, *osterix*:NTR, *osterix*:NTR x *mmp9*:NTR sibling zebrafish and *osterix*:CreERT2mCherry zebrafish, respectively. Fluorophore view images in high brightness and contrast settings (asterisks) to reveal dying cells in NTR+ zebrafish (same adjustments for *osterix*:CreERT2mCherry). White arrowhead, reduced regenerate length, black arrowheads, segment joints. (B) Quantification of experiment shown in (A). One-way Anova (Tukey), excluding *osterix*:CreERT2 (non-sibling). (C) IHC sections of NFP and BrdU-treated 5 dpa *osterix*:mCherry and *osterix*:NTR x *mmp9*:NTR zebrafish fin regenerates. Arrowhead, BrdU+ cells in distal regenerate. (D) Quantification of experiment shown in (C). Kruskal-Wallis. dist., 0-250 µm from regenerate tip, prox., 250 µm from regenerate tip to amp plane. (E) Recovery treatment regimen. (F) 5 and 7 dpa fin regenerates of 3 to 5 dpa NFP treated fin regenerates of *osterix*:NTR x *mmp9*:NTR zebrafish (DMSO treatment during recovery). (G) Increase in regenerate length in *mmp9*:NTR, *osterix*:NTR, *osterix*:NTR x *mmp9*:NTR and *osterix*:mCherry zebrafish during 2 day recovery (DMSO treatments). Dunnett’s T3, excluding *osterix*:mCherry (non-sibling). (H) DMSO and SU5402 recovery treated 7 dpa fin regenerates of *osterix*:NTR x *mmp9*:NTR and *osterix*:mCherry zebrafish. Scale bars (A, F, H) 200 µm, (C) 20 µm.

We wondered how ablation affected proliferation in the regenerate and characterized the consequences of *osterix*+ and *mmp9*+ cell ablation by applying the S-phase marker Bromodeoxyuridine (BrdU) during the last 6 hours of ablation. 5 dpa double ablated fin regenerates showed a normal rate of proliferation in the epidermis; however, reduced mesenchyme (blastema and osteoblast) proliferation (**Fig. 6C, D**), pointing to reduced cell expansion as one cause for impaired regeneration, in addition to death of ablated cells.

Next, we asked whether Fgf signaling would be pivotal in the recovery function of the distal regenerate after impairment of regeneration. To test this, we ablated *mmp9*+ and *osterix*+ cells from 3 to 5 dpa, determined regenerate length as well as *osterix*+ and *osterix*-domain sizes and treated the zebrafish with SU5402 during the recovery phase thereafter (**Fig. 6E**). An increase in regenerate length from 5 to 7 dpa was observed in control conditions without ablation, individual *mmp9* / *osterix* cell ablation, as well as after double ablation (**Fig. 6F, G, Suppl Fig. 10A, B)**; however, addition of regenerative tissue in double ablated fins lagged somewhat behind compared to single cell-type ablated fins (**Fig. 6G, Suppl Fig. 10B**). Notably, Fgf inhibition did not affect recovery in any of the ablation conditions (**Fig. 6H**, **Suppl Fig. 10A, B**) indicating that Fgf signaling is dispensable for recovery function.

### osterix+ osteoblast ablation impairs bone matrix formation and patterning while mmp9+ cell ablation does not

We went on to test whether cell ablation would irreversibly affect specific domains in the elongating regenerate. In order to distinguish recovering from non-recovering regions after ablation, we used the *runx2*:GFP+ reporter as an indicator for (pre-) osteoblast presence in *runx2*:GFP x *osterix*:NTR transgenic zebrafish at 5 dpa (i.e. after 2 days of NTR-mediated *osterix*+ cell ablation), and at 7 dpa after 2 days of recovery (**Fig. 7A-C**). At 5 dpa, *osterix*+ osteoblast-ablated fin regenerates were significantly shorter overall, while the length of individual domains (*runx2*:GFP+ region proximal to fin ray bifurcations, *runx2*:GFP+ region distal to fin ray bifurcations, tip region devoid of *runx2*:GFP signal) was only mildly reduced (**Fig. 7B**). Two recovery days later, the distal *runx2*:GFP+ region showed normal domain length, while the *runx2*:GFP+ region proximal to bifurcation was significantly shorter and of the same length as directly after ablation. In comparison, control treated fins possessed a more than twofold longer pre-bifurcation proximal domain (**Fig. 7A, C**). Furthermore, the size of the tip domain was slightly (albeit insignificantly) longer in fins that had undergone *osterix*+ osteoblast ablation. This indicates that distal regenerate regions less affected by *osterix*+ cell ablation regrow to pre-amputation size, independent of regeneration defects in proximal parts of the fin. It also suggests that distal regenerate cells might partly compensate for impaired regeneration of proximal fin tissue.

**Fig. 7.**
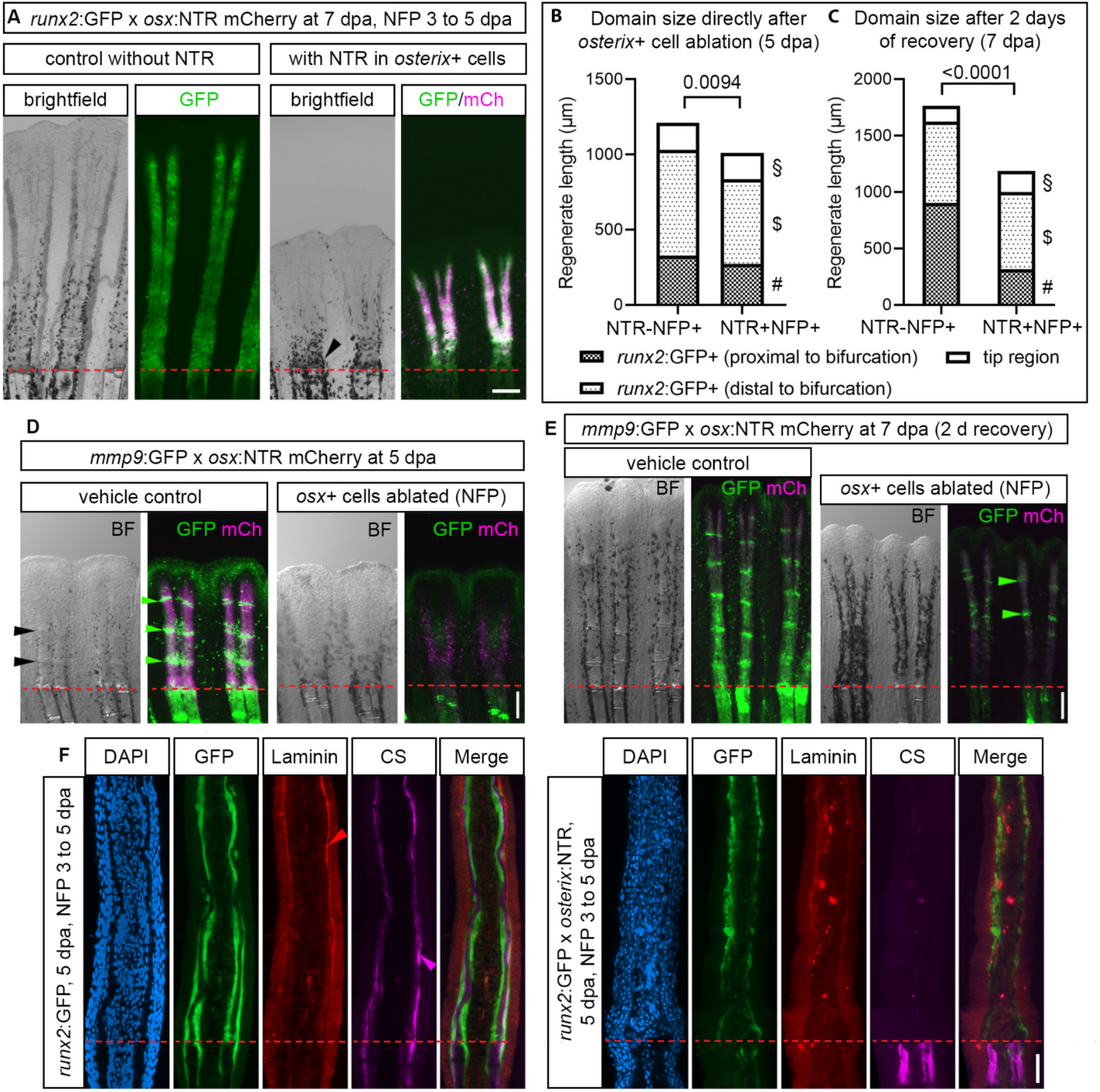
Structural integrity and establishment of segment joints are impaired after *osterix*+ cell ablation. (A) 7 dpa fin regenerates of 3 to 5 dpa NFP treated *runx2*:GFP x *osterix*:NTR zebrafish. Arrowhead, accumulated melanocytes. (B) Regenerate length and domain sizes (proximal vs distal to bifurcation, *runx2*:GFP negative tip region) at 5 dpa after NFP treatment. Welch’s t-tests, Mann-Whitney test for tip region. p(§, tip region)=0.7732, p($, dist. *runx2*:GFP)=0.2564, p(#, prox. *runx2*:GFP)=0.6341. (C) Regenerate length and domain sizes at 7 dpa, after 2d recovery. Welch’s t-tests. p(§, tip region)=0.0624, p($, dist. *runx2*:GFP)=0.7599, p(#, prox. *runx2*:GFP)=0.0100. (D) 5 dpa fin regenerates of 3 to 5 dpa DMSO and NFP treated *mmp9*:GFP x *osterix*:NTR zebrafish. Black and green arrowheads, segment joints. (E) 7 dpa fin regenerates of 3 to 5 dpa DMSO and NFP treated *mmp9*:GFP x *osterix*:NTR zebrafish (2 d recovery). Arrowheads, segment joints. (F) 5 dpa anti-chondroitin sulfate (CS) and anti-Laminin stained *runx2*:GFP x *osterix*:NTR fin regenerate sections after NFP treatment. Arrowheads, Laminin and CS signal in non-ablated fins. Scale bars (A, D, E) 200 µm, (F) 50 µm.

Next, we tested how *mmp9*:GFP+ cells, which partly reside at developing segment joints would react to *osterix*+ cell ablation. In control treatment conditions, *mmp9*:GFP+ cells were visible in up to 3 forming segment joints as well as in the distal portion of the regenerate of *mmp9*:GFP x *osterix*:NTR reporter zebrafish (5 dpa, **Fig. 7D**). In contrast, NFP treated fin regenerates undergoing *osterix*+ cell ablation lacked confined expression of *mmp9*:GFP at segment joints and showed much weaker *mmp9* expression in a single broader domain at about 50 % proximo-distal level of the shorter fin regenerates, potentially reflecting dying cells (**Fig. 7D**). At the same time, the segment joint indicator *pthlha* was not expressed in its usual pattern anymore (**Suppl Fig. 11A**). Inspection of *osterix*+ cell ablated fins showed that segment joints did not form, although control treated fins formed a minimum of one segment joint in a more proximal position (**Fig. 7D, Suppl Fig. 11B**). Notably, in the reverse scenario of *mmp9*+ cell ablation, segment joint formation was observed (black arrowheads in **Fig. 6A, Suppl Fig. 11C**). This indicates pronounced patterning defects in case *osterix*+ osteoblasts are depleted from fin regenerates. We let *mmp9*:GFP x *osterix*:NTR regenerating zebrafish recover from vehicle/NFP treatment for 2 days. During this time, NFP-treated fins re-initiated segment joint formation in the distal part of the regenerate, resulting in an overall reduced segment joint number (**Fig. 7E, Suppl Fig. 11D)**, again indicating independence of the distal regenerate domain of proximal deterioration.

We asked why regenerate outgrowth and patterning were severely affected after *osterix*+ cell ablation and hypothesized that bone matrix secretion and maturation were diminished, leading to a lack of structural support in fin regenerates. We stained for the presence of chondroitin sulfate (CS), an indicator of matrix maturation ^40^ and investigated the presence of Laminin as a measure of basement membrane integrity of the BLWE overlying *osterix*+ osteoblasts ^41^. Ablation of *osterix*+ osteoblasts led to strong CS reduction in the regenerate while CS levels in stump bone matrix were unaffected (**Fig. 7F**). Laminin expression was maintained after ablation; however, showed irregular and ill-defined distribution indicating impaired BLWE integrity after ablation (**Fig. 7F**), which indicates that structural ECM integrity is required for appropriate patterning of the different functional domains in the fin regenerate.

## Discussion

With this work, we provide a dataset on the transcriptomic landscape of rare blastemal, osteoblast and epidermal cell populations of the 3 dpa zebrafish fin regenerate browsable at the Single Cell Portal (https://singlecell.broadinstitute.org/) (reviewer access details needed) which represents a rich resource for researchers investigating appendage regeneration in teleost fish and other species ^42–44^. The dataset contains cells which may have been underrepresented in whole fin regenerate sequencing datasets due to the high prevalence of other cells ^11, 12^ but which are important to investigate the lineage specification of bone forming cells and DMB cells serving as a signaling center throughout regeneration. As we have used cells from fin regenerates after repeated amputation, it can also be used to study potential changes arising after recurrent injury, although our analysis suggests that gene expression in the distal regenerate is generally not altered between different rounds of amputation.

### Spatial arrangement and diversity of blastema cells, BLWE cells and osteoblasts in the regenerate

Research on the origin of skeletal cells in the regenerating appendage elucidated the role of stump tissues and cellular plasticity in the regeneration process ^4–8, 42^. Other studies, such as by Brown et al. (2009), Stewart et al. (2014) and Wehner et al. (2014) have provided insight on the hierarchy and regulation of distal *Runx2*+ pre-osteoblasts, and a more proximal *osterix*+ osteoblast cell pool in the regenerate. Here, we uncovered the complex tissue constitution of the distal regenerate by identifying 5 osteoblast cell populations, including previously described committed osteoblasts, for which we present novel marker genes, two populations of proliferating osteoblasts, osteoblasts with a transcriptomic profile akin to blastema cells, and joint cells located at the segment boundaries. We have analyzed a comparatively high number of osteoblasts (2321 cells) of different maturity and proliferative capacity, and - as we suggest, proximo-distal position. This proximo-distal position (**Fig. 2J**) can be inferred from combinatorial gene expression analysis. Importantly, we identified a population of blastema cells (Blastema3) and a population of osteoblasts (Osteo1), which form a transition zone between blastema and osteoblast cell clusters of the regenerate. The observation that BLWE cell clusters separated completely from blastema and osteoblast cell clusters suggests that lateral BLWE do not contribute to the mesenchymal cell pool (and *vice versa*).

### Committed osteoblasts contribute to bone formation in the proximal regenerate and are supported by immature cells located at the tip

Here, we used genetic lineage tracing and ablation of *osterix*+ and *mmp9*+ cells, including Osteo4, Osteo1 and Osteo2 joint cells, to estimate the contribution of both cell populations to bone formation in the caudal fin regenerate. From 3 to 5 dpa, amplification of Osteo4 cells led to progeny covering ∼80 % of the regenerate’s length demonstrating a considerable contribution, supported by a strong reduction in regenerate length after *osterix*+ cell ablation during regeneration. At the same time, the restricted distal expansion of Osteo4 progeny suggested a somewhat limited renewal capacity, which is reflected by the comparatively low number of *pcna*+ expressing cells in this population. In salamanders, transplantation experiments of different Cre-labeled populations (*Col1a2* vs *Prrx1*) suggested that distal skeletal elements of the regenerated limb are derived from non-skeletal connective tissue cells^42^. In our experiments, lineage tracing showed the absence of Osteo4 progeny from the distal 250 µm of the regenerate, suggesting alternative sources for this region. Candidate populations are *stmn1a*+ Osteo3 osteoblasts which have a high proliferative activity (**Suppl Fig. 4G)**, and Osteo1 cells, which are found at the transition zone between blastemal and osteoblast cell clusters in our sc RNA sequencing dataset. Notably, strong *mmp9* expression is found in cells belonging to these and the Osteo0 clusters (**Suppl Fig. 12**), in addition to previously reported expression of *mmp9* in segment joint cells ^7^. Occasional expansion of *mmp9*:CreERT2-converted cell clones at the very tip of the regenerate (**Fig. 3E, H**) suggests contribution of *mmp9*+ cells to an osteoblast progenitor cell pool in this distal region of the regenerate. This is supported by the observed effects on simultaneous *mmp9*+ and *osterix*+ cell ablation, which decreased fin regenerate growth more dramatically than individual ablation.

The origin of osteoblast progenitors in the distal portion of the regenerate has been vague. Distal Runx2+ cells are maintained as a pre-osteoblast cell pool under the influence of Wnt signaling^10^. In line with this, we identified a region of *runx2*:GFP+ pre-osteoblasts showing very weak GFP fluorescence in close proximity to the DMB. The respective cells often co-labeled with mCherry protein produced in *siam*:mCherry Wnt-responsive cells (**Suppl Fig. 8C**). We suggest that these cells have just started to differentiate into pre-osteoblasts and that they are derived from blastema cells. Alternatively, these cells may represent descendants of *runx2*:GFP+ osteoblasts that have lost *gfp* expression and have relocated distally. A second population of more proximal mCherry, GFP double+ cells may partly be explained by overlap of their respective transcript domains. However, no such zone of concomitant *gfp*/*mCherry* expression was detected in double transgenic *osterix*:GFP x *siam*:mCherry zebrafish, in agreement with low Wnt signaling in *osterix*+ osteoblasts ^10, 25^. Nevertheless, a considerable amount of GFP/mCherry protein double+ cells were detected in the proximal region of the regenerate. We explain this discrepancy between mRNA and protein levels with a relocation of distal blastema cells to a more proximal position, or by a distal movement of osteoblasts towards the tip, resulting in mixing of cells. This is in line with our TEM data showing a gradual phenotype change of cells underlying the BLWE along the proximodistal direction. We conclude that a lineage dependence exists between blastema cells and osteoblasts in the distal fin regenerate and that this transition is laid out by Blastema3 and Osteo1 cells, followed by proliferative Osteo0/3 cells. *mfap5*, *stmn1a*, *twist2* and *runx2*:GFP are markers for this transition. Since blastema clusters were continuous in our analysis, actinotrichia-forming cells and even DMB cells may ultimately be involved in this process. Interestingly, photoconversion of DMB kaede-labelled cells, carried out by Wehner et al. (2014) led to a small population of converted cells localizing to a slightly proximal position [Fig. S2I in ^25^]. Altogether, we propose a model in which committed osteoblasts maintain themselves in the proximal domain of the regenerate, while the distal domain of the regenerate is populated by immature osteoblast progenitors derived from *mmp9*+ pre-osteoblasts, and *runx2*:GFP+ cells that are provided by blastema cells. Notably, this distal domain is able to compensate for diminished regenerate growth after a challenge such as Fgf signaling inhibition, demonstrating the adaptability of this region.

### The activity and size of the different regenerate domains depends on the presence of growth factors and reflects the need for continued regenerate growth

Here, we show that manipulation of growth factor signaling affects domain extension in the regenerate, similar to altered domain sizes after osteoblast ablation. Inhibition of Fgf signaling specifically reduced the domain size of *osterix*+ cells while leaving their reporter activity as well as the size of the *osterix*-, *siam*+ distal tip region unchanged during inhibition. This indicates a profound effect of Fgf signaling on newly differentiating *osterix*+ cells. This effect is reversible and both the size and activity of the *siam*+ *osterix*-tip region and the activity of osteoblasts in the *osterix*:nGFP+ domain increase after discontinuation of Fgf suppression. Recovery of fin regenerate length is then completed within a few days, which is accompanied by the presence of an enlarged tip domain, suggesting intrinsic mechanisms detecting the need for enhanced regeneration which is likely carried out by enhanced proliferation at the junction of distal pre-osteoblasts and *osterix*+ osteoblasts. Notably, spatial information cannot be overridden by Fgf suppression. The same is true for fins that have undergone a more dramatic, simultaneous ablation of both *mmp9*+ and *osterix*+ cells in fin regenerates. These fins catch up in distal growth when ablation is stopped, albeit at a somewhat slower pace than fins with single *mmp9*+ or *osterix*+ cell ablation (**Fig. 6G**).

### Regenerate domains compensate for each other as long as structural support is provided

Here, we show that after ablation of *osterix*+ cells in the proximal regenerate, the *runx2*:GFP+ cell population distal to ray bifurcation expands normally, and that the *runx2*-tip region remained slightly longer in *osterix*+ cell ablated fins than in control-treated fins. In contrast, the domain proximal to ray bifurcations, which did lose structural support i.e. bone matrix, did not recover, therefore leading to a significantly reduced domain length. In the future, it will be important to test whether *osterix*+ cell ablated fins can grow to their pre-amputation size thanks to the tip region or whether regenerates remain significantly shorter because of the lack of proximal tissue. Given that the *runx2*:GFP-tip region is already slightly longer after a 2-day recovery, this region might take over ‘responsibility’ for regenerate growth. This would be in agreement with other experimental settings we have tested, in which the tip region of the regenerate increased in size to make up for suppressed regenerate growth.

In our work, we detected patterning defects affecting segment joint formation in case *osterix*+ osteoblasts lining segments (but not segment boundaries) were ablated. To our surprise, we did not see segment joint formation defects in case of *mmp9*+ cell (i.e. joint forming cell) ablation. This suggests that *osterix*+ cells and the resulting deposited bone matrix is required for appropriate segmentation of the fin. Altogether, this demonstrates a pivotal function of committed osteoblasts, but also pre-osteoblasts and blastema cells for growth, bone formation and patterning of the regenerating vertebrate appendage.

## Supporting information

Supplemental Table 1 reg1 vs rest deg

Supplemental Table 2 DEGS main clusters

Supplemental Table 3 Blastema GO terms

Supplemental Table 4 Osteo GO terms

Supplemental Table 5 Basal GO terms

Supplemental Table 6 DEGs Blastema clusters

Supplemental Table 7 DEGs Osteo clusters

Supplemental Table 8 DEGs Basal clusters

## Acknowledgements

This work was supported by the DFG TRR67 and SPP2084 µBone (grants to F.K.). A.D., S.Re. and M.L. are supported via the NGS Competence Centre of the DFG (GZ: INST 269/768-1, “Förderinitiative Hochdurchsatz-Sequenzierung: DRESDEN-concept Genome Center”). This project has received funding from the European Research Council (ERC) under the European Union’s Horizon 2020 research and innovation program (grant agreement no. 950349). This work was supported by the DRESDEN-concept Genome Center, CMCB Flow Cytometry Core and Light Microscopy Facilities of the TU Dresden CMCB Technology Platform. The work at the TU Dresden is co-financed with tax revenues based on the budget agreed by the Saxonian Landtag. We are grateful to M.-A. Akimenko, D. Wehner, R. Bernitz, G. Weidinger, S. Hans and A. Kawakami for sharing reagents and transgenic zebrafish. We thank K. Geurtzen, A. Pollack, and L. Pohnert for technical assistance, L. Limant for help with measurements, and M. Fischer, J. Michling, D. Mögel and S. Kunadt for excellent fish care. Many thanks to D. Wehner for useful comments and H. Knopf for proof-reading the manuscript.

## Author contributions

Conceptualization, F.K., N.C., and A.C.L.D.; Methodology, F.K., N.C., A.C.L.D., F.R., and S.R.; Formal Analysis, N.C., A.C.L.D., F.R., F.K., T.K. and M.L.; Investigation, N.C., A.C.L.D., F.R., F.K., T.K., S.R.; Resources: A.D., S.R., F.K.; Writing - Original Draft, F.K., N.C., and A.C.L.D.; Writing - Review & Editing: F.K., N.C., A.C.L.D., F.R., T.K., and S.R.; Visualization, N.C., A.C.L.D., F.K. and F.R.; Supervision, F.K. and S.R., Funding Acquisition: F.K., A.D., and S.R.

## Methods

### Animal experiments

All procedures were approved by and performed according to the animal handling and research regulations of the Landesdirektion Dresden (Permit numbers: AZ 24D-9168.11-1/2008-1, AZ 24-9168.11-1/2011-52, AZ DD24.1-5131/354/87, AZ DD24.1-5131/450/4, AZ 25-

5131/496/56 and respective amendments).

### Transgenic zebrafish and husbandry

The transgenic fish lines used in this study have been described: *siam*:mCherry=Tg(7xTCF-Xla.Sia:NLS-mCherry) ^47^, *shh*:GFP=Tg(−2.7shha:GFP) ^48^, *osterix:CreERT2-p2a-mCherry=*Tg(Ola.Sp7:CreERT2-P2A-mCherry)tud8, *runx2*:GFP=Tg(Hsa.RUNX2-Mmu.Fos:EGFP)zf259 ^4^, *osterix*:mCherry=Tg(Ola.Sp7:mCherry)zf131, *osterix*:GFP=Tg(Ola.Sp7:NLS-GFP)zf132 ^49^, *mmp9*:GFP=TgBAC(mmp9:EGFP)tyt201, *mmp9*:NTR=TgBAC(mmp9:EGFP-NTR)tyt207, *mmp9*:CreERT2=TgBAC(mmp9:Cre-ERT2, cryaa:EGFP)tyt208 ^7^, dsRed2GFP=Tg(hsp70l:loxP-DsRed2-loxP-nlsEGFP)tud9 ^4^, *osterix*:NTR=Tg(Ola.Sp7:mCherry-Eco.NfsB)pd46 ^38^. The artificial promoter *siam* drives mCherry expression in Wnt responsive cells in the DMB and some more proximal cells ^47^. The *shha* promoter element ^48^ activates expression of GFP in BLWE cells. Osteoblasts of different maturity were labeled by the activity of *osterix* and *RUNX2* promoter elements ^4^. Fish were bred and maintained as described ^4^.

### Fin clips

Fin clips were performed as described ^50^. 3 dpa regenerates were obtained from fin-clipped zebrafish. Fish were allowed to regenerate in 28°C water.

### Tissue dissociation and flow cytometry

Fin regenerates of quadruple transgenic fish (*siam:mCherry, shh:GFP, osterix:CreERT2-p2a-mCherry, runx2:GFP)* were harvested at 3 dpa, cut into small pieces with a scalpel and transferred into 1 ml collagenase-dispase solution (1 mg/ml in PBS, Roche #10269638001) for 10 min at 28°C. The sample was pipetted slowly up and down with an elongated, flame polished Pasteur pipette. The procedure was repeated 4 times with decreasing inner tip diameters of the Pasteur pipettes until a homogenous solution was obtained. The dissociates were poured onto an equilibrated 70 µm cell strainer and collected in 10 ml HBSS solution (without CaCl_2_ and MgCl_2_, Gibco #12082739). After centrifugation (15 minutes, 1800 rpm, 4°C), the supernatant was discarded and the remaining cell pellet resuspended in 500 µl 2% BSA in PBS. Calcein violet (1µl 10mM, Invitrogen #C34858) was added to the cell solution and incubated for 30 minutes. Calcein violet, GFP and mCherry+ cells were collected in 50 µl 2% BSA in PBS via fluorescence-activated cell sorting (BD LSR Fortessa) and processed for single-cell RNA sequencing analysis based on 10X Genomics (10X Chromium system, 10X library preparation according to the manufacturer’s instructions).

### Single-cell RNA sequencing analysis

For each experiment about 8000 cells were flow-sorted into BSA-coated PCR tubes containing 1 µl of PBS with 0.04 % BSA. All cells were carefully mixed with reverse transcription mix before loading them in a Chromium Single Cell A Chip on the 10X Genomics Chromium controller ^51^ and processed further following the guidelines of the 10X Genomics user manual for single cell 3’ RNA-seq v2. In short, the droplets were directly subjected to reverse transcription, the emulsion was broken and cDNA was purified using silane beads. After amplification of cDNA with 12 cycles, it underwent a purification with 0.6 volume of SPRI select beads. After quality check and quantification using the Fragment Analyzer (Agilent), 30 ng cDNA were used to prepare sc RNA-seq libraries - involving fragmentation, dA-Tailing, adapter ligation and a 12 cycles indexing PCR based on manufacturer’s guidelines. After quantification, both libraries were sequenced on an Illumina Nextseq500 system in paired-end mode with 26 bp/57 bp (for read 1 and 2 respectively), thus generating ∼60-90 mio. fragments for the transcriptome library on average.

A custom reference based on GRCz10, Ensembl annotation e98 was created, by first adding the sequences of *gfp* and *mCherry* as separate chromosomes to the fa file and the gtf file and then building the cellranger reference using cellranger mkref. Next, fastq files were processed with cellranger count from 10X genomics (https://support.10xgenomics.com/single-cell-gene-expression/software/pipelines/latest/using/count) version 3.0.0 using the custom reference. This resulted in a dataset with 2532, 3601, 2481 and 3413 mean counts per cell and median number of 300, 532, 975 and 508 detected genes per cell, and 7028, 4707, 2846 and 4535 cells for regeneration experiments Reg1, Reg2, Reg3 and Reg4, respectively.

For the downstream analysis of the 10X data, current best practices were followed ^13^. The filtered gene counts matrices were read with scanpy 1.6.0 ^52^. Next, cells were filtered based on the number of total counts, the number of detected genes and with sample specific thresholds: Only cells with more than 2000, 1000, 1000 and 1000 total counts as well as more than 600, 300, 300 and 400 detected genes for samples Reg1, Reg2, Reg3 and Reg4, respectively, were kept for the analysis. Furthermore, cells with more than 5% of mitochondrial reads were filtered. After quality control, our dataset consisted of 1342, 1410, 2102 and 1814 cells from the respective regeneration experiment. Only genes which were detected in more than 3 cells (counting all samples together) were kept for downstream analysis. Normalization was performed with the scanpy function sc.pp.normalize_total and the data was log-transformed with sc.pp.log1p. Highly variable genes were detected with sc.pp.highly_variable_genes setting n_top_genes=4000. Principal component analysis was performed on the highly variable genes. A neighbor graph was constructed with sc.pp.neighbors setting n_neighbors=30 and n_pcs=15. Next, a UMAP was constructed with sc.tl.umap setting min_dist=0.9 ^53^. Clustering was done using sc.tl.leiden with resolution=0.2. Marker genes were computed with sc.tl.rank_genes_groups. Two clusters were identified as blood vessels and immune cells (autofluorescent cells) and those were excluded from further analysis, leaving 1219 (Reg1), 1331 (Reg2), 2007 (Reg3) and 1705 cells (Reg4). Next, principal component analysis, neighborhood graph construction, UMAP computation and marker gene detection were repeated with the same functions and parameters as above. In result, the presented UMAPs show 691 BLWE cells (Basal), 3250 blastema cells (Blastema), and 2321 osteoblasts (Osteo). The 3 main clusters were sub-clustered using sc.tl.leiden setting the restrict_to parameter to the respective cluster. The resolution parameter was set to 0.25, 0.1 and 0.35 for the blastema, BLWE and osteoblast clusters, respectively. For each sub-cluster, marker genes were computed as above but using only the cells of the respective main cluster as a reference. Trajectory inference was performed with sc.tl.paga and the paga plot was created sc.pl.paga_compare setting threshold=0.1 ^14^. For differential expression analysis, we used a limma-voom workflow: only highly expressed genes were considered, i.e. genes with a cpm value>1 in more than 25% of the cells ^54^. The AnnData object was converted to an edgeR DGEList object using anndata2ri (https://github.com/theislab/anndata2ri) and scran ^55, 56^. Normalisation factors of the raw count matrix were computed with edgeR’s calcNormFactors function. The number of detected genes per cell was scaled to zero mean and unit variance and added as a co-factor to the design formula. The other factor that was added was the condition (Reg1 vs Reg2-4 pooled together). Next, the DGE list was transformed using limma’s voom function ^57^. Next, edgeR’s functions lmFit, contrasts.fit, treat (with lfc=log2(1.5)) and topTreat were used to generate a list of differentially expressed genes. Next, pseudospace coordinates were computed. For the lateral coordinate, the gene set *lamb1a*, *wnt5b*, *shha* and *phlda2* were used. A laterality score was computed using the scanpy function sc.tl.score_genes. Next, the transcriptome data object was subset to the lateral gene set. Diffusion pseudotime was computed by running sc.pp.neighbors with n_neighbors=50, sc.tl.diffmap and sc.tl.dpt on the subsetted transcriptome data. For sc.tl.dpt, the cell with the highest laterality score was used as the root. The lateral coordinate of pseudospace was then computed by a rank transformation of the pseudotime. For the distal coordinate, the same computation was performed using the gene set *aldh1a2*, *wnt5a*, *fgf3*, *fgf10a*, *igf2b*, *msx3*, *msx1b*, *cdh4*, *dkk1b*, *wnt3a*, *dkk1a*, *dlx5a*, *junba*, *junbb*, *msx2b*, *spry4* and *wnt10a*.

The dataset is available for the reviewers to access at the Single Cell Portal:

**Accession:** currently for reviewers

**URL:** currently for reviewers

**PIN:** currently for reviewers

### Statistical analysis

Statistical analysis was performed using Prism 6 (GraphPad Software, La Jolla, CA, USA), with the statistical tests and corresponding p values reported in the figures and respective legends. All values represent the mean ± SD unless otherwise stated.

### RNA *in situ* hybridization and histology

The plasmids to obtain probes for *fgf24* (restriction digest with NotI, transcription with T7), and *pthlha* (restriction digest with BamHI, transcription with SP6) have been published ^30, 58^. *mCherry* (restriction digest with HindIII and transcription with T7), and *gfp* (restriction digest with EcoRI, transcription with T3) plasmids were provided by Gilbert Weidinger. The *CreERT2* probe plasmid (restriction digest with AgeI, transcription with T7) was provided by Stefan Hans. The *tnc in situ* probe plasmid was generated by René Bernitz and provided by Daniel Wehner. The *pthlha in situ* probe plasmid ^30^ was provided by Marie-Andrée Akimenko. Additional probes have been synthesized from pCR-Blunt II-TOPO vectors after cloning of the respective sequences by using the Zero Blunt TOPO PCR cloning Kit (Invitrogen) according to the manufacturer’s instructions. The forward and reverse oligonucleotides to amplify the cDNA sequences are listed below, along with the corresponding fragment sizes.

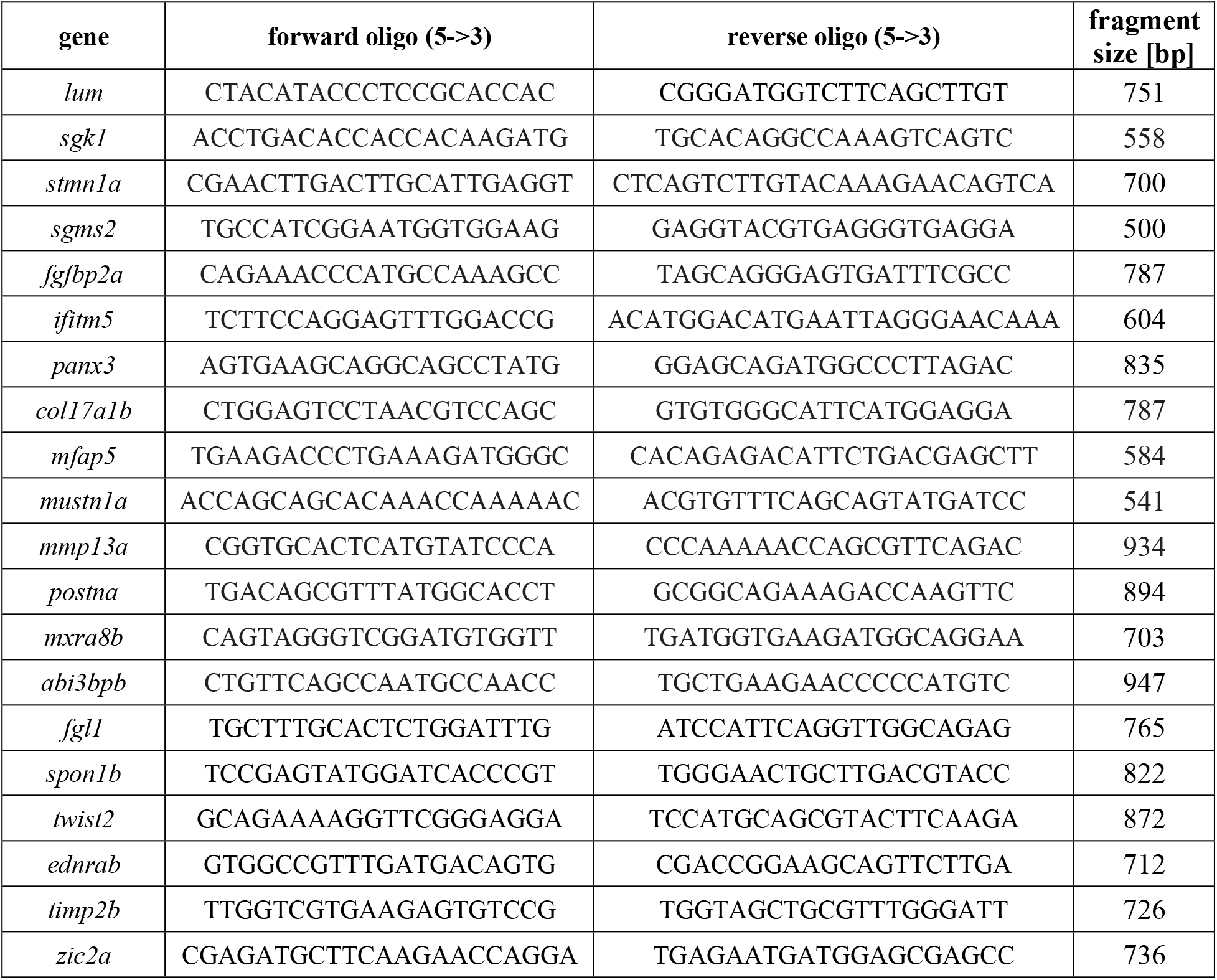
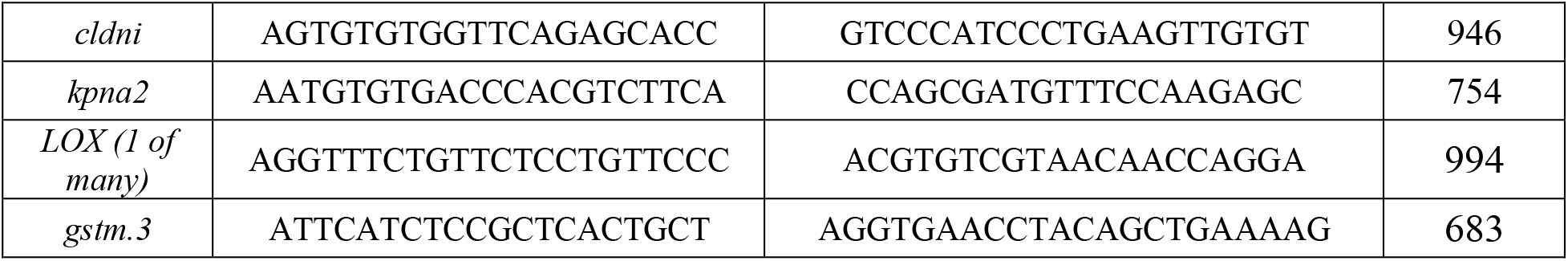

RNA *in situ* hybridization and histological processing of whole mount fin regenerates was performed as previously described ^59^. For double RNA ISH, NBT/BCIP staining (stock solution Roche, 11681451001) was combined with Fast Red staining for simultaneous detection of two transcripts. After staining with NBT/BCIP, fins were briefly rinsed twice in PBT (PBS, 0.1% Tween 20) and transferred to new tubes. This was followed by incubation in 0.1M Glycin/HCL, pH 2,2 + 0.1% Tween (2x 5 minutes) and washing with PBT (4x 5 minutes). 200µl Anti-Fluorescein-AP, Fab fragments antibody (Roche, 11426339810) was added (1:2000 dilution in PBT + 2 mg/ml BSA + 2% sheep serum) and incubated over night at 4°C. The fins were kept dark and washed in PBT (2x 5 minutes, 6x 30 minutes). Fins were then washed in 0.1M Tris pH 8.2 + 0.1% Tween (3x 5 minutes). SIGMAFAST™ Fast Red TR/Napthol AS-MX Tablets (Sigma, F4648) were dissolved according to the manufacturer’s instructions. Specimens were kept dark and at RT in 500 µl Fast Red staining solution until the staining had developed. The staining reaction was stopped by two brief washes with PBT and one wash in STOP solution (46 ml 0.5M NaH_2_PO_4_ + 4 ml 0.5M NaHPO_4_ pH 5.8 + 1mM EDTA + 0,1% Tween 20). Fins were stored in 80% glycerol/20% STOP solution thereafter and processed for cryosectioning. Cryosections were imaged with a Zeiss 10x/0.45 Plan-Apochromat air objective on an ApoTome1 equipped with a Zeiss AxioCam MRc color CCD camera and a Zeiss ZEN blue (v 2012) software.

### Immunohistochemistry

Preparation of tissue for cryosectioning and immunofluorescence were performed as described^4^. 12 µm cryosections were obtained with a Cryostat HM560. Primary antibodies used were: chicken anti-GFP (Abcam, ab13970) at 1:2000, mouse anti-mCherry (Clontech, 632543) at 1:450, rabbit anti-dsRed (Clontech, 632496) at 1:300, mouse anti-chondroitin sulfate (Sigma, C8035) at 1:300, rabbit anti-Laminin (Sigma, L9393) at 1:200, rat anti-BrdU (Novus Bio, NB500-169) at 1:300. Secondary antibodies used were: goat anti-chicken-Alexa 488 (Thermo Fisher, A-11039), goat anti-mouse-Alexa 555 (Thermo Fisher, A-21424), goat anti-mouse-Alexa 633 (Thermo Fisher, A-21046), goat anti-rat-Alexa 633 (Thermo Fisher, A-21094) at 1:1000.

### Combined RNA *in situ* hybridization and immunohistochemistry

RNA *in situ* hybridization was performed as indicated previously but after incubation of anti-DIG-AP, primary antibodies (chicken anti-GFP, Abcam, ab13970, 1:2000 and rabbit anti-DsRed, Clontech, 632496, 1:200) were incubated overnight at 4°C. Fins were washed in PBT and incubated with secondary antibodies (goat anti-chicken-Alexa 488, Thermo Fisher, A-11039 and goat anti-rabbit-Alexa 633, Thermo Fisher, A-21071) at 1:500. After immunohistochemistry, fins were stained with Fast Red solution and cryosectioned. Cryosections were imaged with a Zeiss 20x/0.8 Plan-Apochromat air objective on a LSM 980/MP inverse equipped with a Zeiss Zen2 blue (v 3.0.79.00004 HF4) software.

### Live imaging, quantification of fluorescent cells and plot profile measurements

For quantification of GFP expression fish were anesthetized with 0.02% Tricaine (MS222) and their fins were imaged with a Zeiss SteREO Discovery.V12 microscope equipped with a AxioCam MRm camera and AxioVison software version 4.7.1.0. Identical settings for magnification, exposure time, gain, and contrast were used. Live imaging of *siam*:mCherry x *runx2*:GFP and *siam*:mCherry x *osterix*:GFP transgenic zebrafish was performed on a Dragonfly Spinning Disk confocal equipped with an Andor Zyla PLUS monochrome sCMOS camera and Fusion software with a z interval of 2 µm and a 30x silicone objective. GFP and mCherry double+ cells were counted in 50 µm intervals beginning with the most distal mCherry signal (0 µm) up to 400 µm. Quantification of reporter fluorescence along fin rays of transgenic *siam*:mCherry x *osterix*:GFP zebrafish was done with the plot profile tool in Image J/Fiji.

### Fate mapping

Transgenic fish (*osterix*:CreERT2-p2a-mCherry x *hsp70l*:R2nlsGFP; *mmp9*:CreERT2, *cryaa*:EGFP x *hsp70l*:R2nlsGFP and *osterix*:CreERT2-p2a-mCherry, *mmp9*:CreERT2, *cryaa:EGFP* x *hsp70l*:R2nlsGFP x *Ola.Actb*:LOXP-DsRed2LOXP-EGFP) ^4, 7, 60^ were injected intraperitoneally with 10µl 2.5mM 4-hydroxytamoxifen (Sigma #H7904) in PBS or with the vehicle control ethanol in PBS at 2.5 dpa (approx. 36 hpa). The fish were heat shocked for 1h at 37°C at 66, 84 and 108 hpa. Fins were imaged at 3 dpa (approx. 72 hpa), 4 dpa (approx. 96 hpa) and 5 dpa (approx. 120 hpa) and analyzed for recombination events.

### Electron microscopy

Fin regenerates were cut off distal to the amputation site and were fixed in 4% PFA in 100 mM phosphate buffer, pH 7.4. Samples were further dissected for embedding into epoxy resin and processed according to a modified protocol for serial block face SEM ^61^ using osmium tetroxide (OsO_4_), thiocarbohydrazide (TCH), and again OsO_4_ to generate enhanced membrane contrast ^62, 63^. In brief, samples were postfixed overnight in modified Karnovsky fixative (2% glutaraldehyde/2% formaldehyde in 50 mM HEPES, pH 7.4), followed by post-fixation in a 2% aqueous OsO_4_ solution containing 1.5% potassium ferrocyanide and 2mM CaCl_2_ (30 minutes on ice), washes in water, 1% TCH in water (20 minutes at RT), washes in water and a second osmium contrasting step in 2% OsO_4_/water (30 minutes on ice). Samples were washed and *en-bloc* contrasted with 1% uranyl acetate/water for 2 h on ice, washed in water, and dehydrated in a graded series of ethanol/water mixtures (30%, 50%, 70%, 90%, 96%), followed by 3 changes in pure ethanol on molecular sieve. Samples were infiltrated into the epon substitute EMBed 812 (resin/ethanol mixtures: 1:3, 1:1, 3:1 for 1h each, followed by pure resin overnight and for 5hrs), embedded into flat embedding molds, and cured at 65°C overnight. Ultrathin sections (70 nm) were prepared with a Leica UC6 ultramicrotome (Leica Microsystems, Wetzlar, Germany) and a diamond knife (Diatome, Nidau, Switzerland), collected on formvar-coated slot grids, and stained with lead citrate ^64^ and uranyl acetate. Mounted sections were analyzed with a JEM 1400Plus transmission electron microscope (JEOL, Freising, Germany) at 80 kV and images were taken with a Ruby digital camera (JEOL).

### Drug treatments

Fgf inhibition was performed with 17 µM SU5402 (Calbiochem) or the vehicle control dimethylsulfoxide (DMSO) in fish water at the times indicated. Solutions were changed daily. 50 mg/ml Bromodeoxyuridine (BrdU) stock solutions were prepared in DMSO and used at 5 mM (Fig. 5A-F, Fig. 7A-F) or 2.5 mM (Fig. 6C-H) in selected experiments, either in fish water, in fish water supplemented with NFP or DMSO, or in fish water supplemented with SU5402 or DMSO. BrdU treatment was performed during the last 6 hours of day 5 post amputation. NTR mediated ablation was performed with 1 µM NFP in fish water. Fin regenerates were either fixed and/or photographed at 5 dpa or zebrafish were allowed to recover from treatment for 2 days before fin regenerates were fixed and/or photographed.

## Supplemental information

### Supplemental figure legends

**Suppl Fig. 1.**
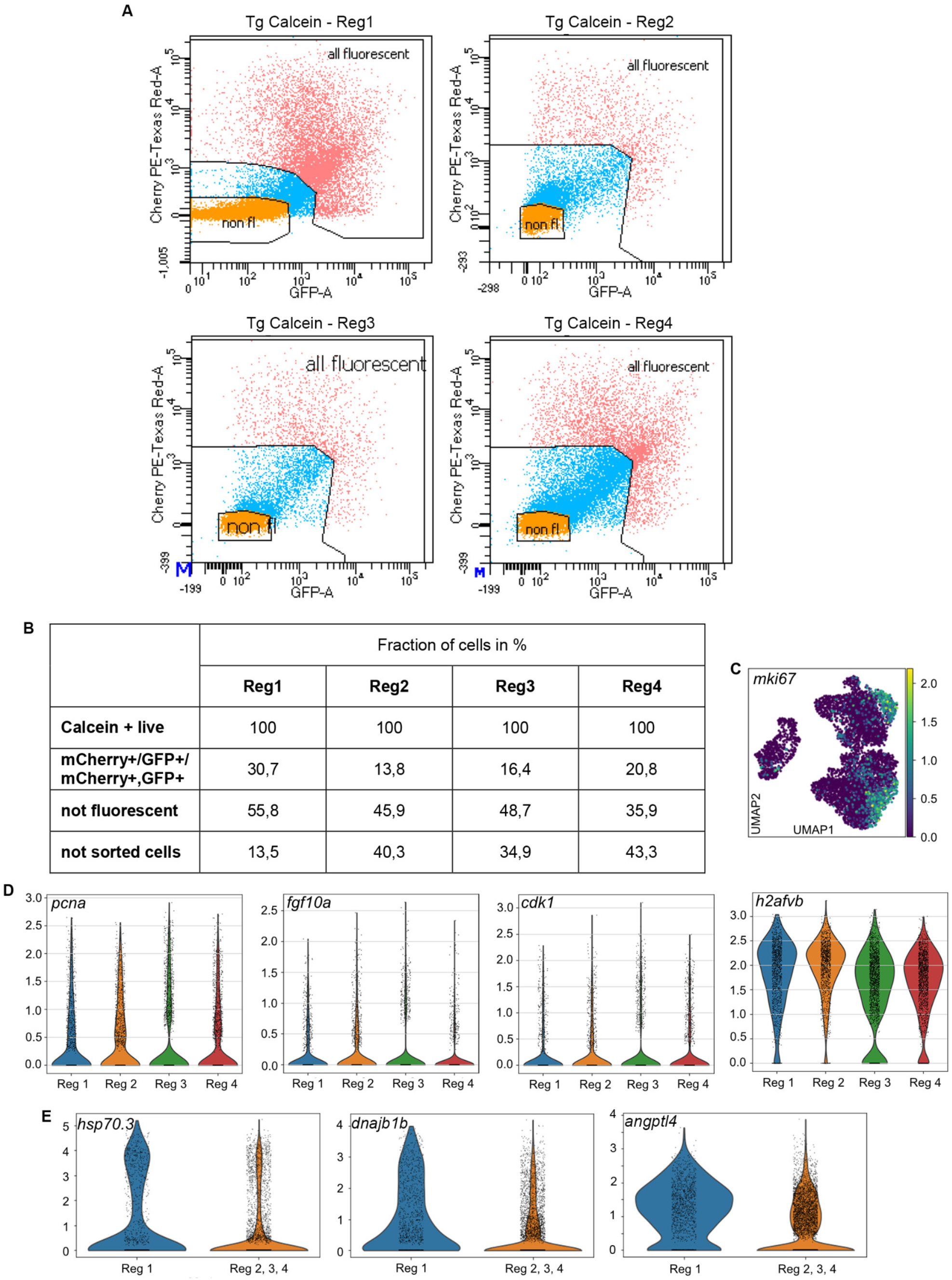
FACS conditions to isolate cells of interest, mki67 UMAP, and comparison of different regeneration experiments. (A) Gatings for Reg1-Reg4. (B) Percentages of fluorescent (mCherry single+, GFP single+, mcherry/GFP double+), non-fluorescent and other non-sorted cells in Reg1-Reg4. (C) UMAP of the proliferation marker *mki67*. (D) Violin plots of *pcna*, *fgf10a*, *cdk1*, and *h2afvb* in different regeneration experiments. (E) Violin plots of *hsp70.3*, *dnajb1b*, and *angptl4* in Reg1 vs Reg2-4.

**Suppl Fig. 2.**
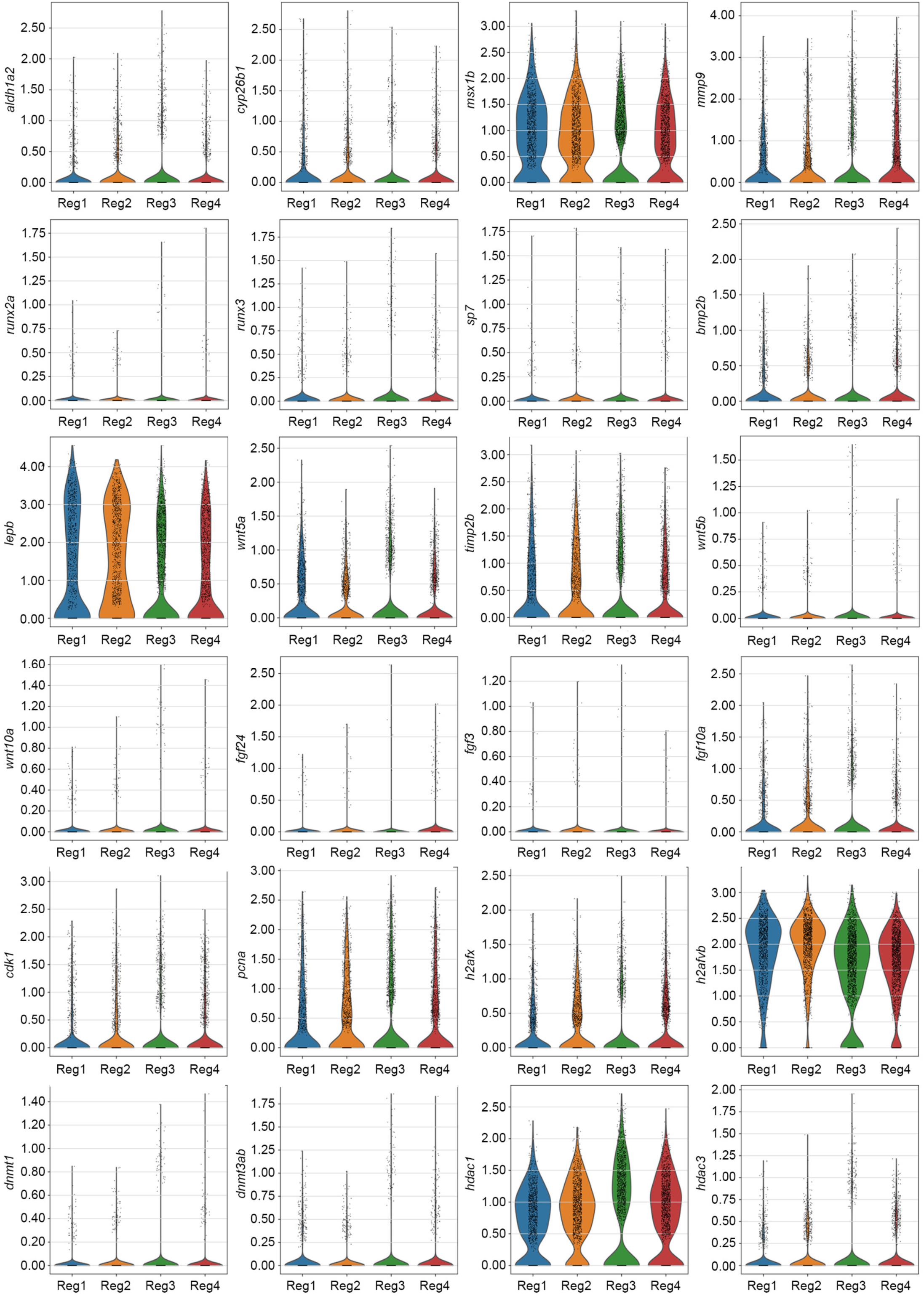
Selected gene expression in Reg1-Reg4 samples. Neither a consistent boost or suppression of gene expression can be detected after repeated amputation. Violin plots of *aldh1a2, cyp26b1, msx1b, mmp9, runx2a, runx3, sp7, bmp2b, lepb, wnt5a, timp2b, wnt5b, wnt10a, fgf24, fgf3, fgf10a, cdk1, pcna, h2afx, h2afvb, dnmt1, dnmt3ab, hdac1, hdac3* in Reg1-4.

**Suppl Fig. 3.**
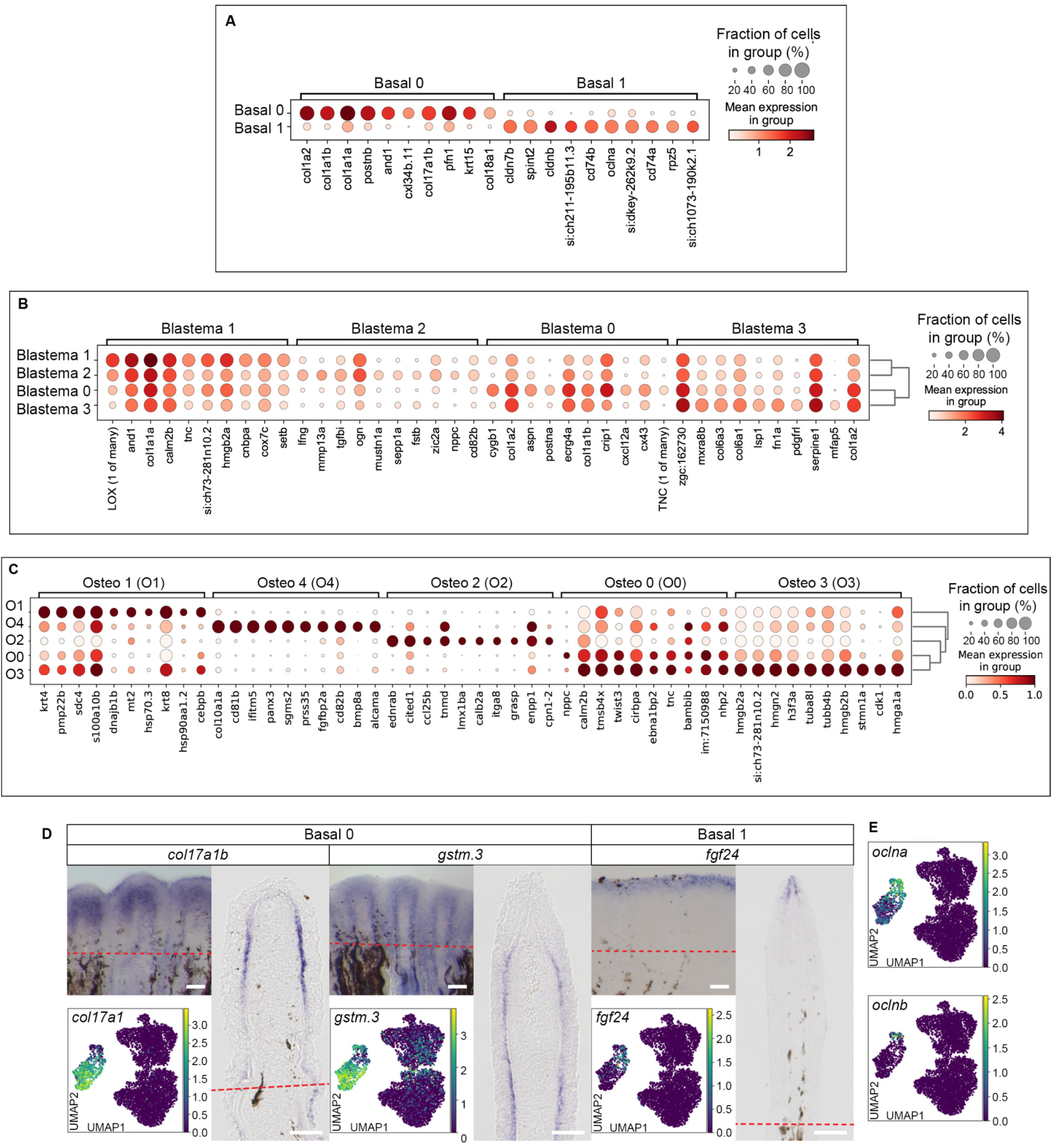
Gene expression in subclusters and discrimination of Basal0 and Basal1 cell clusters. (A) Gene expression in Basal subclusters. (B) Gene expression in Blastema subclusters. (C) Gene expression in Osteo subclusters. (D) Whole mount RNA *in situ* hybridization view, UMAP and cryosection view of *col17a1b*, *gstm.3* and *fgf24* expression. (E) UMAP views of *occludin a (oclna)* and *occludin b (oclnb)* expression.

**Suppl Fig. 4.**
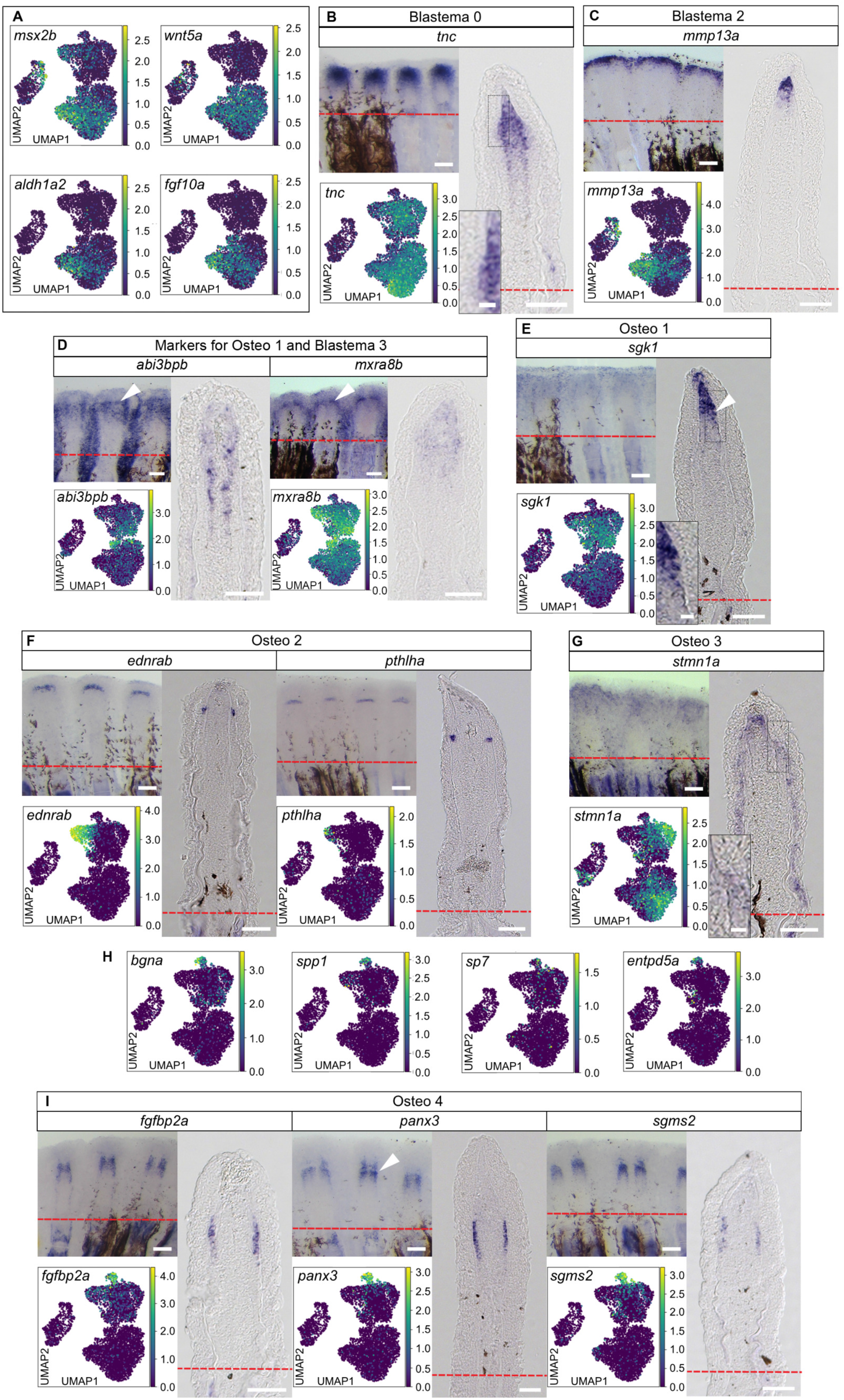
Blastema and osteoblast markers. (A) Expression of known blastema markers *msx2b, wnt5a, aldh1a2, fgf10a* (UMAP views). (B) *tnc* expression in Blastema0. (C) *mmp13a* expression. (D) *abi3bpb* and *mxra8b* expression in Osteo1 and Blastema3 cells. (E) Non-exclusive *sgk1* expression in Osteo1. (F) *ednrab* and *pthlha* expression. (G) Non-exclusive *stmn1a* expression in Osteo3. (H) UMAPs of *bgna*, *spp1*, *sp7* and *entpd5* expression enriched in Osteo4. (I) *fgfbp2a*, *panx3* and *sgms2* expression. (A)-(G), (I) UMAP, WMISH and cryosection views. Scale bars whole mounts 100 µm, cryosections 50 µm, insets 10 µm.

**Suppl Fig. 5.**
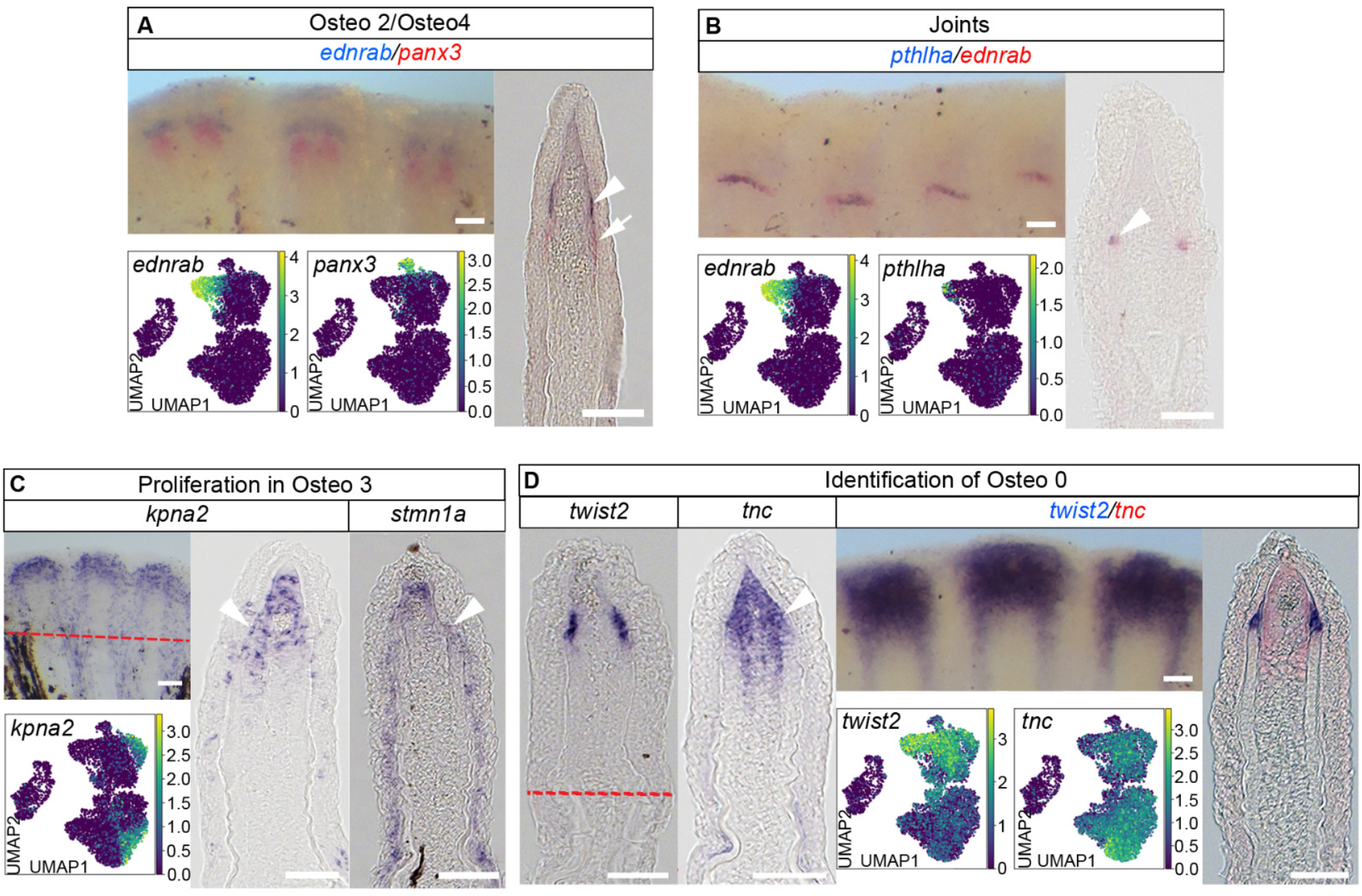
Characterization of specific osteoblast clusters. (A) Distinct locations of Osteo2 (*ednrab*+, arrowhead) and Osteo4 cells (*panx3*+, arrow). Double ISH. (B) Co-labeling of *pthlha* and *ednrab* in Osteo2 cells (arrowhead). Double ISH. (C) Non-exclusive *kpna2* and *stmn1a* expression in Osteo3 (arrowheads). (D) Co-labeling of *tnc* and *twist2* in Osteo0 cells. Individual and double ISH. (A)-(D) UMAP, WMISH and cryosection views. Scale bars whole mounts 100 µm, cryosections 50 µm, insets 10 µm.

**Suppl Fig. 6.**
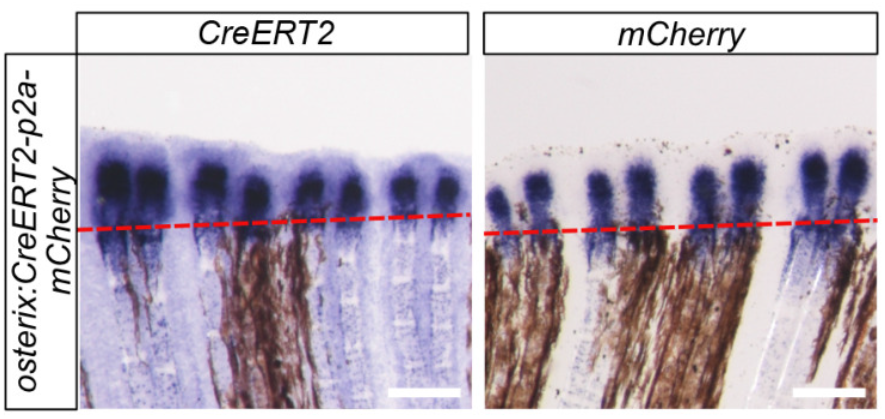
Expression of *CreERT2* and *mCherry* in transgenic *osterix*:CreERT2-p2a-mCherry fins. Expression is broad at 3 dpa. Whole mount ISH. Scale bar 100 µm.

**Suppl Fig. 7.**
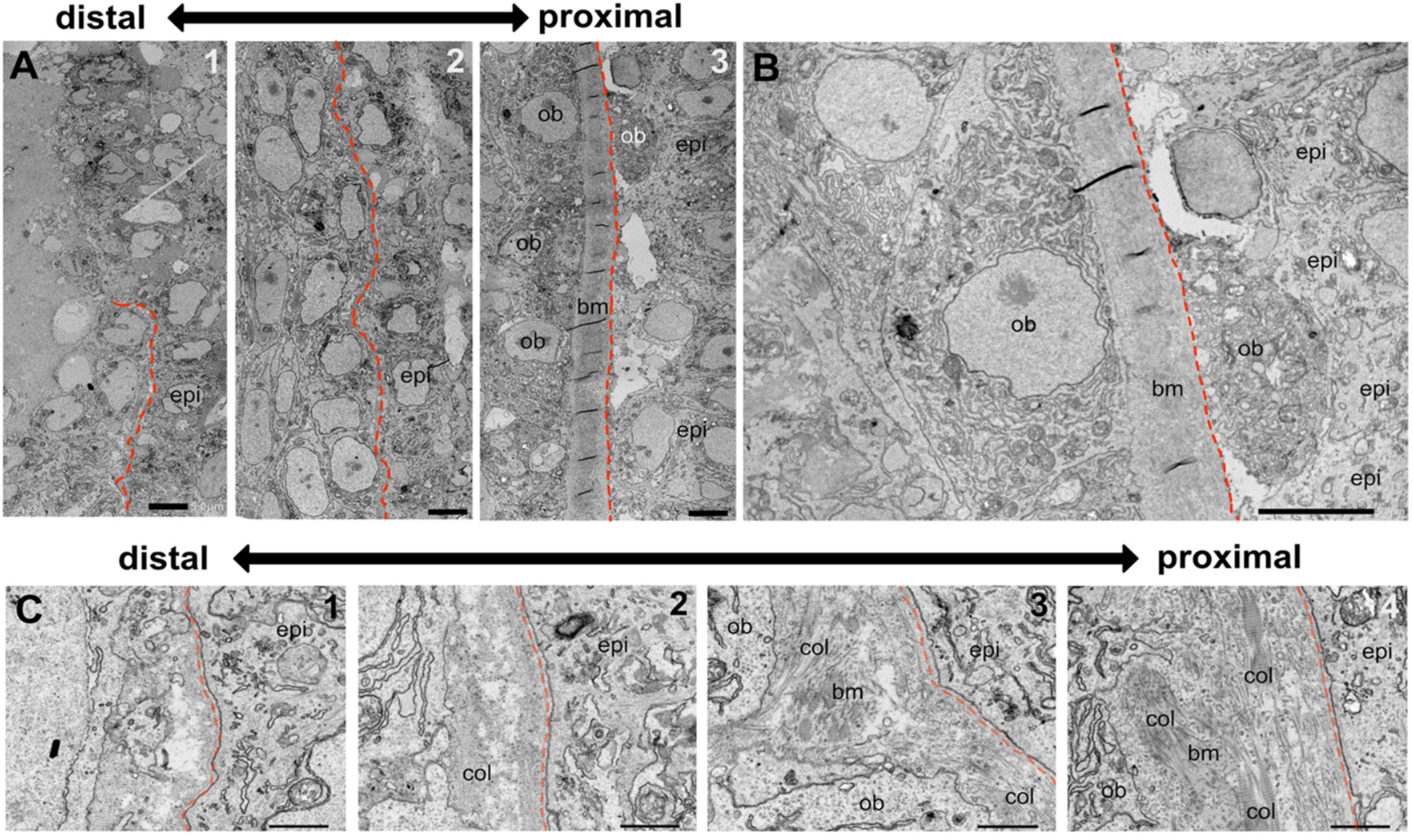
TEM of osteoblasts underlying the BLWE. (A) Sequential overview images from distal (1) via intermediate (2) to proximal (3) regenerate regions. Red dashed lines, subepithelial basal lamina. epi, wound epidermis. Osteoblasts (ob) in more proximal regions begin to produce a thicker ECM layer (bone matrix, bm). (B) ob in A3 at higher magnification. Note the ob beyond the ECM/bm layer below the epi. (C) Images highlighting the ECM layer between epi and ob precursors/osteoblasts from distal to proximal. ECM thickness with collagen fibers (col) and electron dense bm material is more pronounced proximally (3,4). Scale bar (A) 5 µm, (B) 1 µm.

**Suppl Fig. 8.**
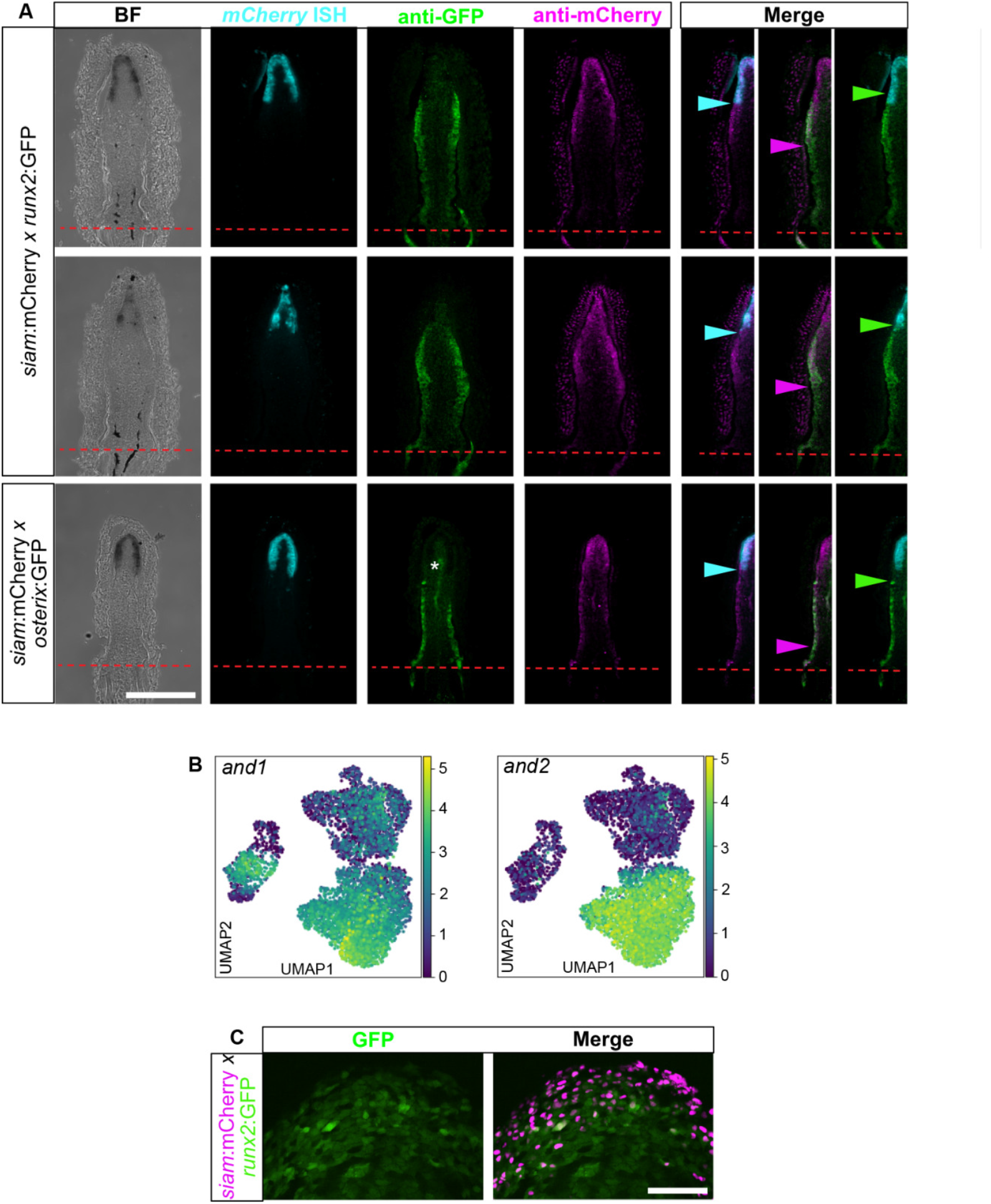
Expression ranges of distal blastema and osteoblast markers differ at the mRNA and protein level. (A) Combined ISH and immunohistochemistry against *mCherry* mRNA, mCherry protein and GFP protein in transgenic *siam*:mCherry x *runx2*:GFP 3 dpa fin regenerates. The *mCherry* mRNA domain is much more restricted than the mCherry protein domain suggesting mixture of mCherry/GFP protein+ cells after transcription of *mCherry* has stopped. Asterisk, autofluorescent endothelial cells, cyan and green arrowheads, proximal limits of *mCherry* RNA and mCherry protein, respectively, magenta arrowhead, distal limit of GFP protein. Scale bar 100 µm. (B) UMAPs of *and1* and *and2* showing broad expression in the blastema. (C) Weak GFP protein expression is detectable in the distal blastema of transgenic *siam*:mCherry x *runx2*:GFP zebrafish at 3 dpa. Scale bar 100 µm.

**Suppl Fig. 9.**
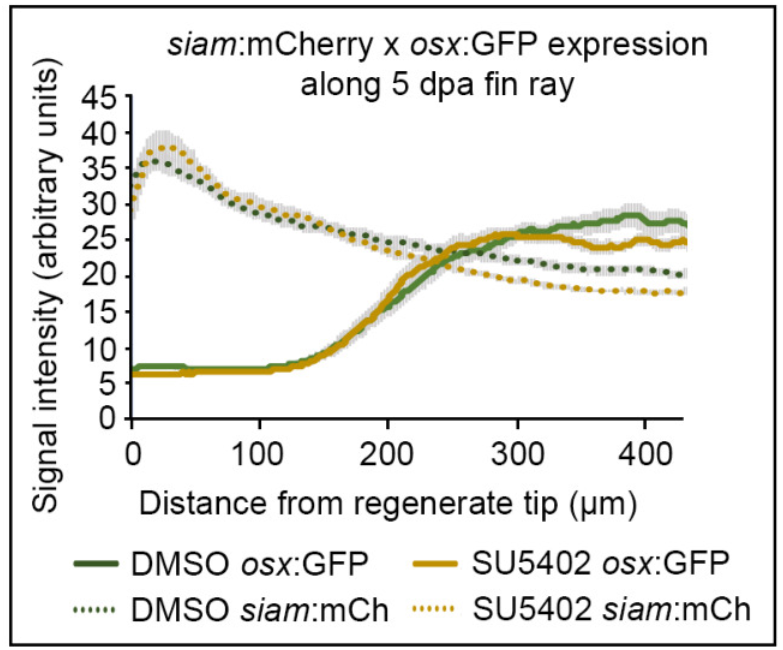
Fluorescence signal intensity of transgenic reporters (*osterix*:GFP, *siam*:mCherry) along the fin regenerate at 5 dpa, treated with DMSO or SU5402 from 3 to 5 dpa. Mean ± SEM.

**Suppl Fig. 10.**
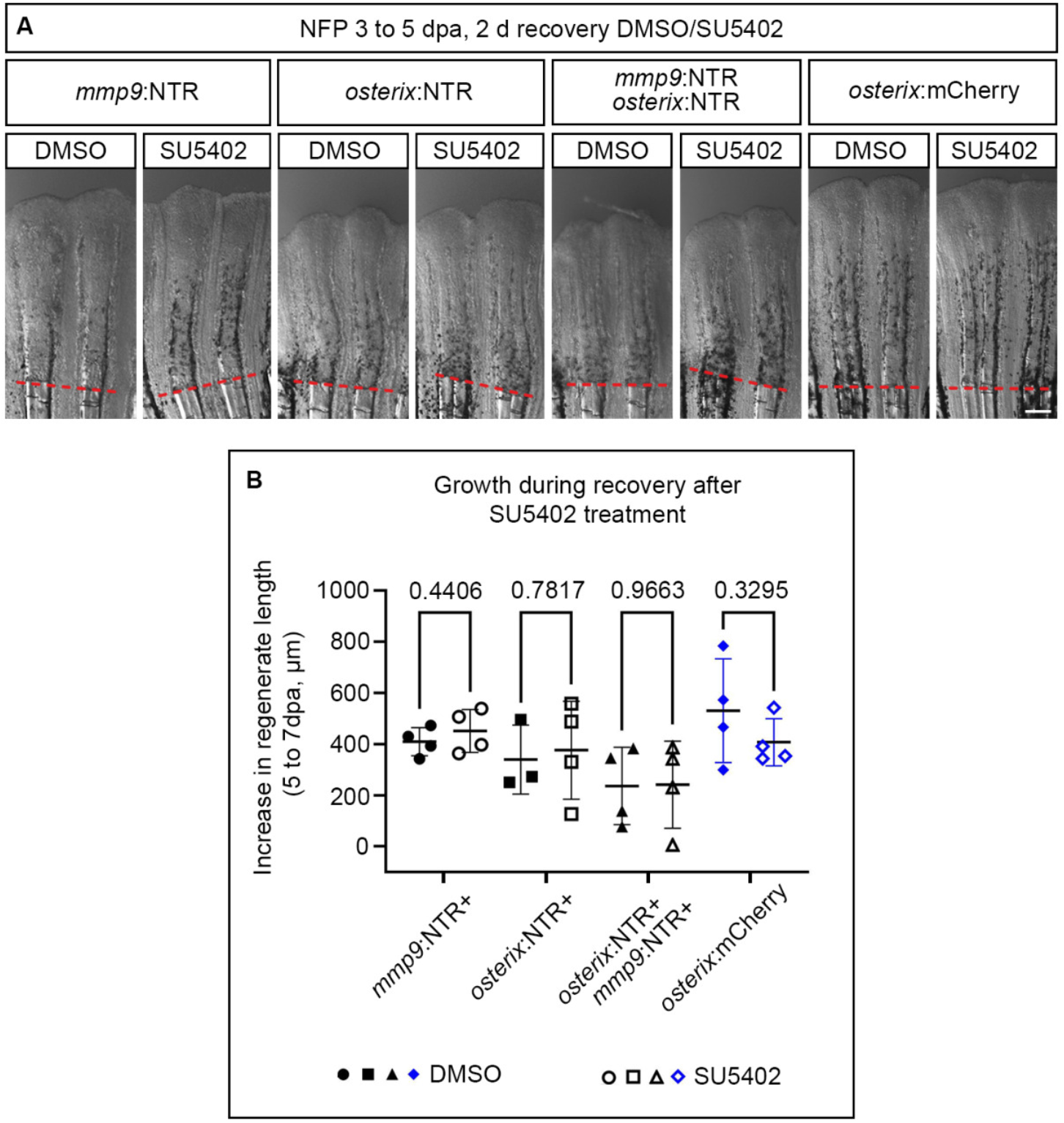
Regenerate recovery is not affected by Fgf inhibition. (A) Representative images of fin regenerates treated either with DMSO or SU5402 during recovery after ablation. Scale bar 200 µm. (B) Quantification of increase of regenerate length during recovery period [experiment shown in (A)]. Welch’s t-tests.

**Suppl Fig. 11.**
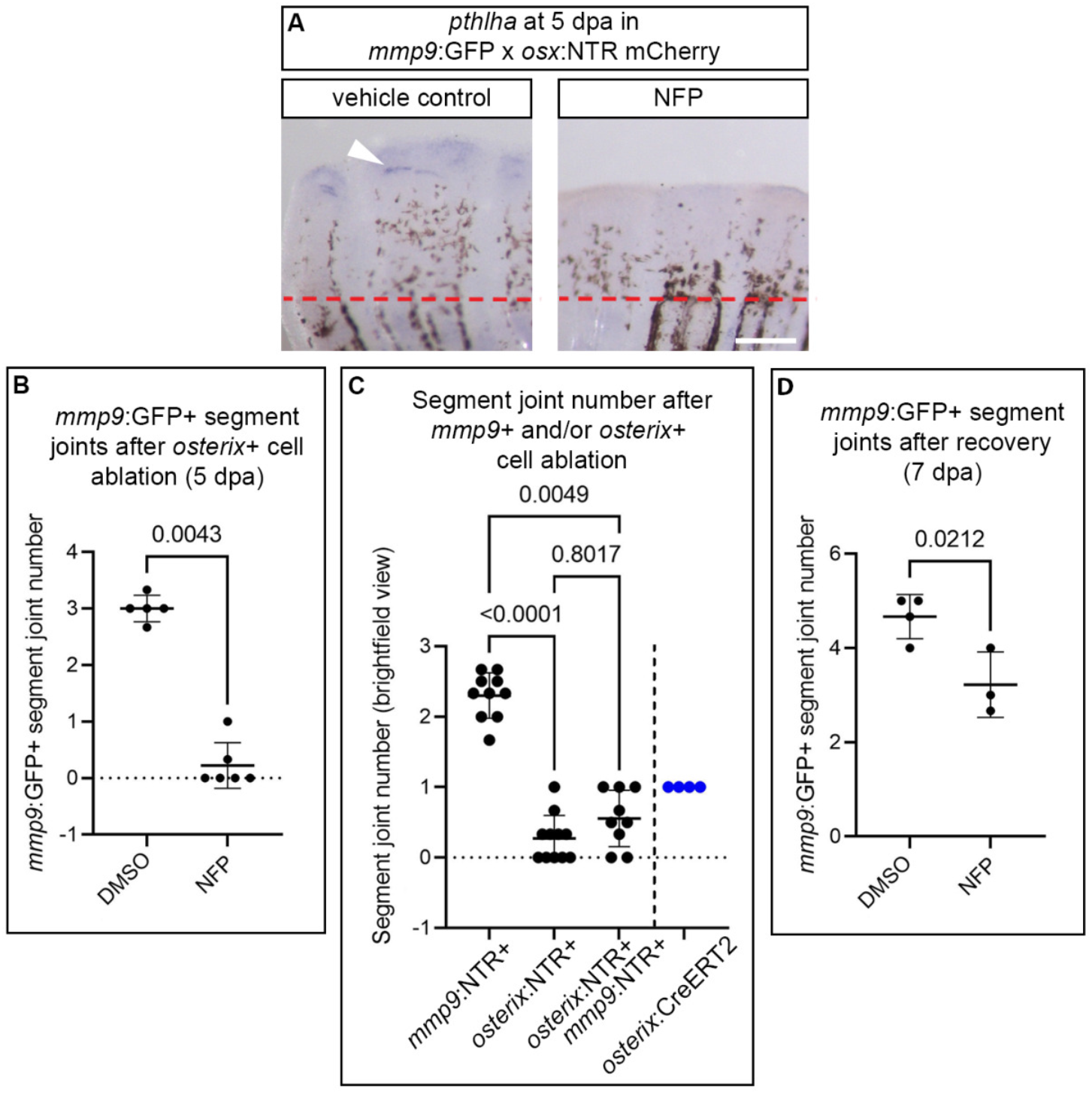
Altered segment joint formation upon *osterix*+ cell ablation. (A) *pthlha* expression in vehicle control (DMSO) treated vs NFP treated *mmp9*:GFP x *osterix*:NTR mCherry treated fin regenerates. Arrowhead pointing at expression at prospective joint forming sites. Scale bar 200 µm. (B) Quantification of *mmp9*:GFP+ segment joint number in vehicle control (DMSO) treated vs NFP *mmp9*:GFP x *osterix*:NTR mCherry treated fin regenerates at 5 dpa (experiment shown in Fig. 7D). Mann-Whitney. (C) Quantification of segment joints visible in the brightfield channel at 5 dpa after either *mmp9*:NTR+ cell ablation, *osterix*:NTR+ cell ablation, combined *mmp9*:NTR+ and *osterix*:NTR+ cell ablation, or control NFP treatment in (non-sibling) *osterix*:CreERT2 zebrafish (experiment shown in Fig. 6A). Kruskal-Wallis. (D) Quantification of *mmp9*:GFP+ segment joint number in vehicle control (DMSO) treated vs NFP *mmp9*:GFP x *osterix*:NTR mCherry treated fin regenerates after recovery at 7 dpa (experiment shown in Fig. 7E). Unpaired two-tailed t-test. (B)-(D) Average segment numbers calculated from 2-3 dorsal fin rays per individual.

**Suppl Fig. 12.**
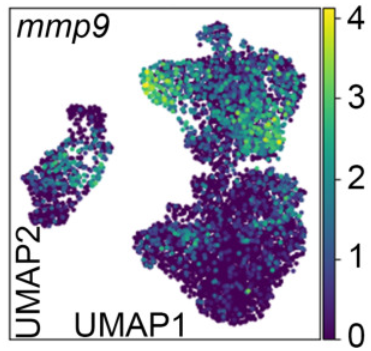
UMAP view of *mmp9* expression. Expression is not restricted to joint cells.

**Suppl Table 9.**
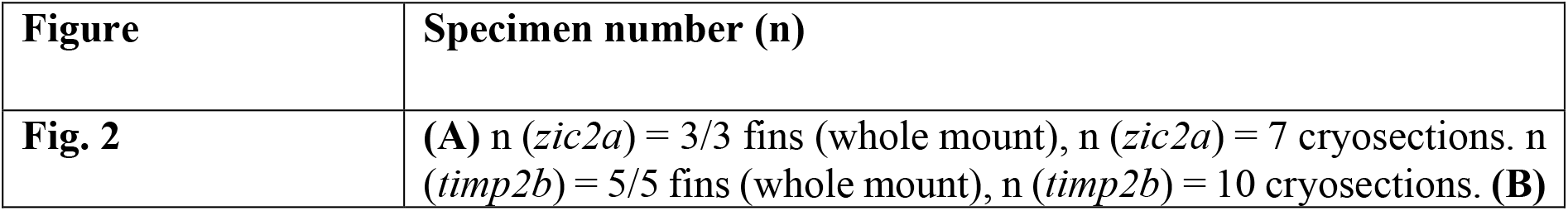
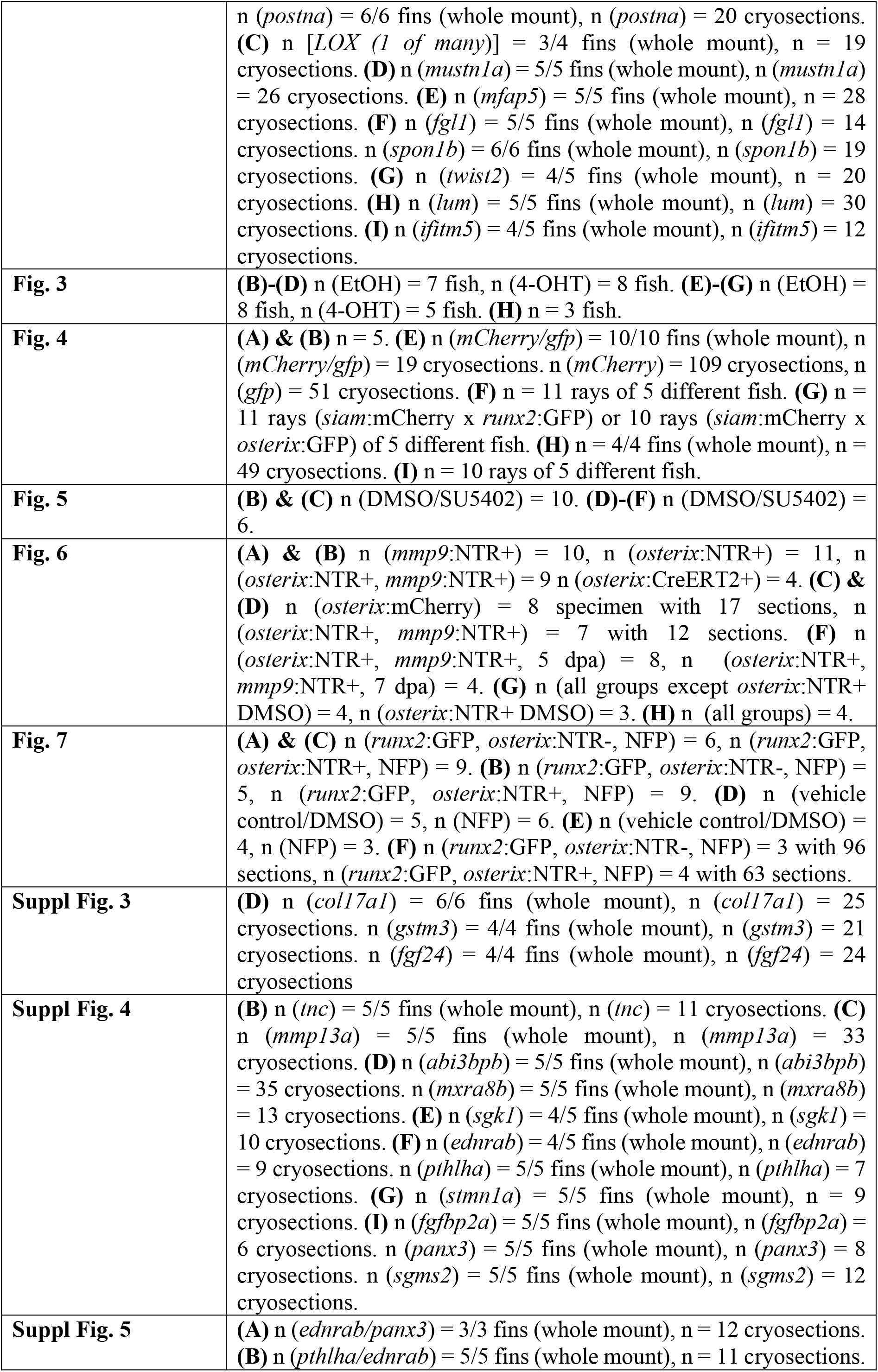
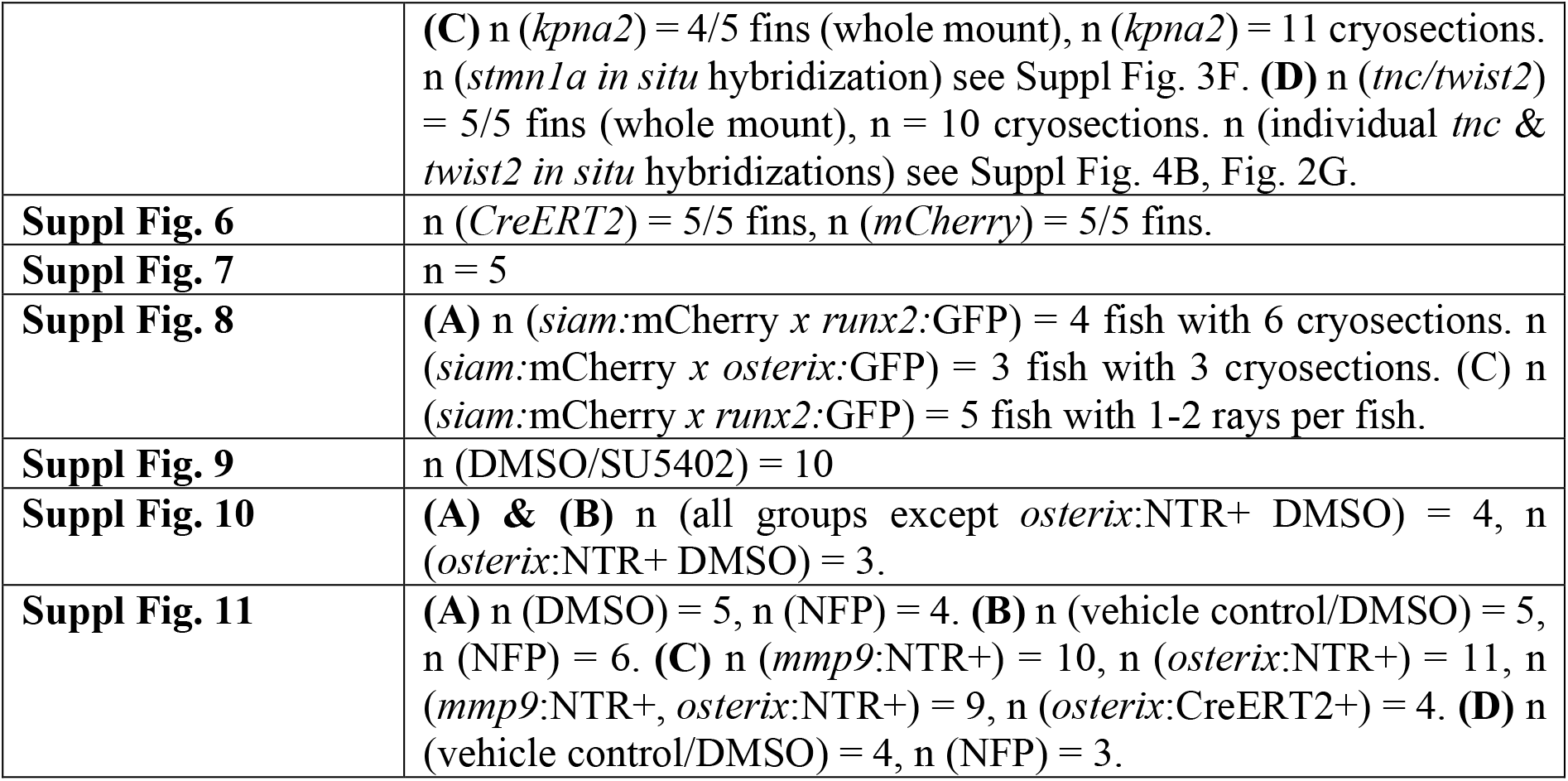
Number of specimens used in experiments. For whole mount RNA ISH: first number = number of specimens displaying the presented staining pattern, second number = total number of specimens used, e.g. 4/5 = 4 out of 5 specimens display the depicted expression pattern.

**Suppl Table 10.**
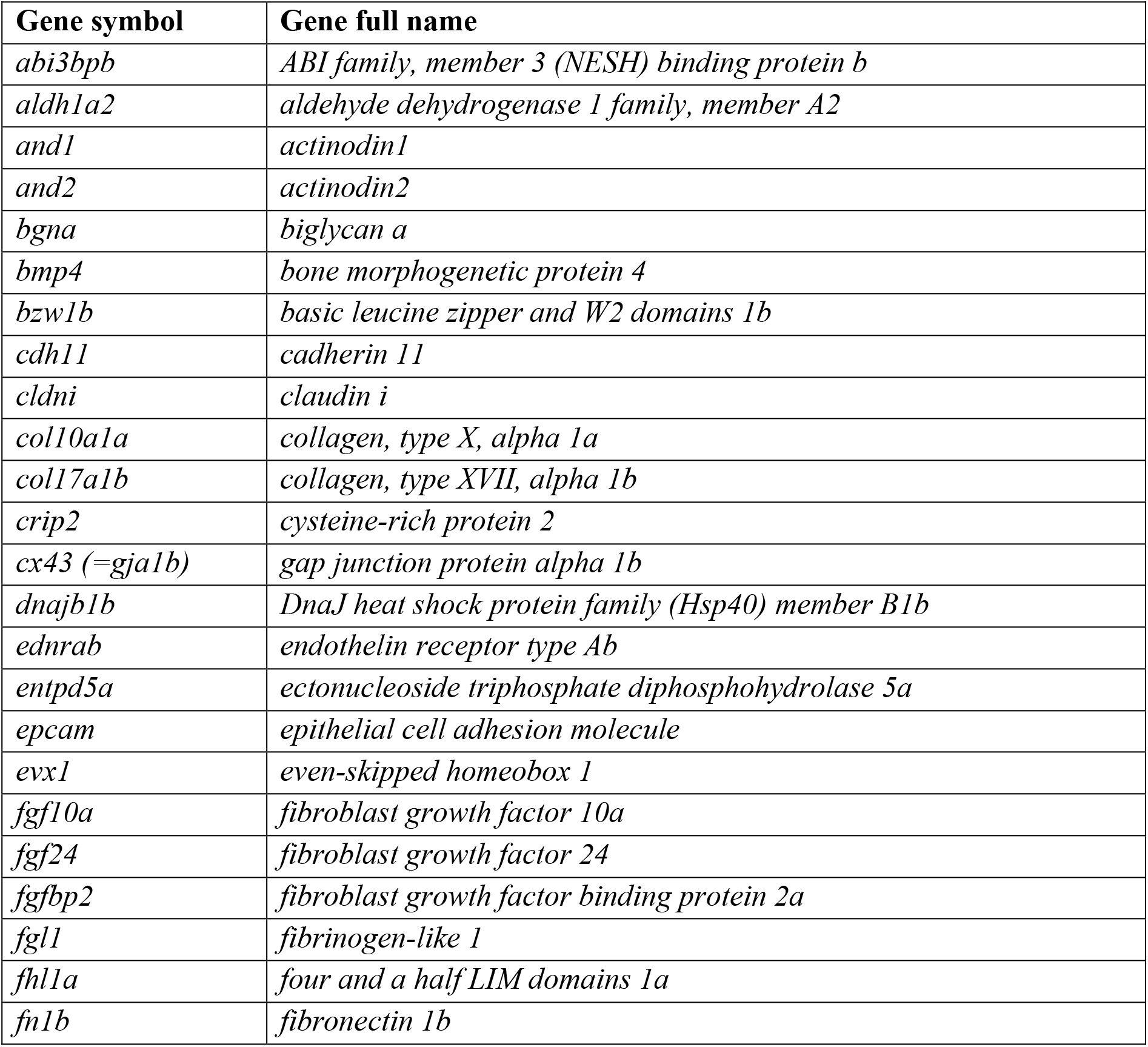
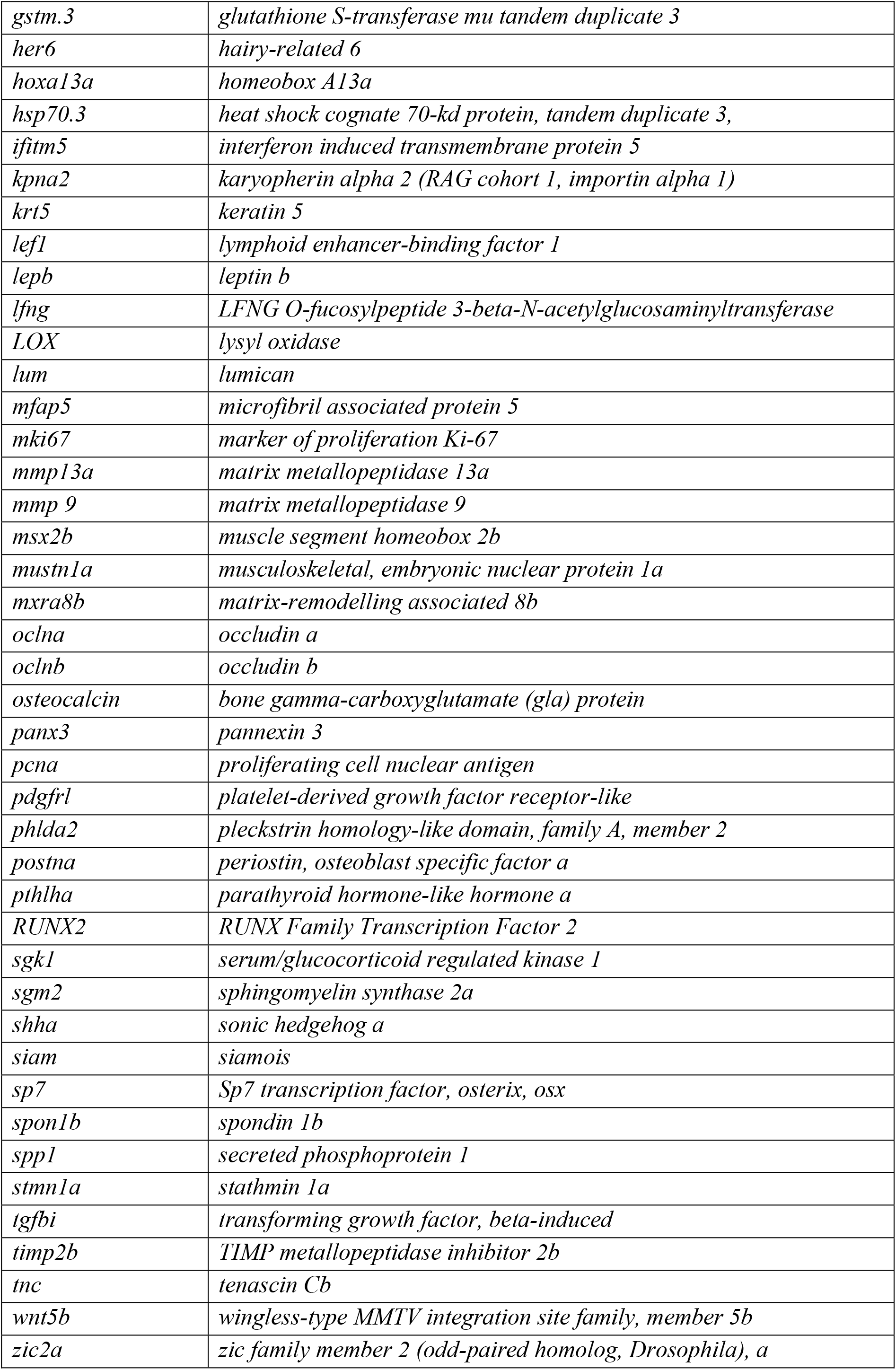
Full gene names in alphabetical order.

## References

1. Dietrich, K., Fiedler, I., Kurzyukova, A., López-Delgado, A.C., McGowan, L., Geurtzen, K., Hammond, C.L., Busse, B., and Knopf, F. (2021). Skeletal biology and disease modeling in zebrafish. J. Bone Miner. Res., jbmr.4256–jbmr.4256. 10.1002/jbmr.4256.

2. Sehring, I.M., and Weidinger, G. (2020). Recent advancements in understanding fin regeneration in zebrafish. WIREs Dev. Biol. 9. 10.1002/wdev.367.

3. Nechiporuk, A., and Keating, M.T. (2002). A proliferation gradient between proximal and msxb-expressing distal blastema directs zebrafish fin regeneration. Development 129, 2607–2617. 10.1242/dev.129.11.2607.

4. Knopf, F., Hammond, C., Chekuru, A., Kurth, T., Hans, S., Weber, C.W., Mahatma, G., Fisher, S., Brand, M., Schulte-Merker, S., et al. (2011). Bone Regenerates via Dedifferentiation of Osteoblasts in the Zebrafish Fin. Dev. Cell 20, 713–724. 10.1016/j.devcel.2011.04.014.

5. Sousa, S., Afonso, N., Bensimon-Brito, A., Fonseca, M., Simões, M., Leon, J., Roehl, H., Cancela, M.L., Jacinto, A., Amsterdam, A., et al. (2011). Differentiated skeletal cells contribute to blastema formation during zebrafish fin regeneration. Development 138, 3897–3905. 10.1242/dev.064717.

6. Stewart, S., and Stankunas, K. (2012). Limited dedifferentiation provides replacement tissue during zebrafish fin regeneration. Dev. Biol. 365, 339–349. 10.1016/j.ydbio.2012.02.031.

7. Ando, K., Shibata, E., Hans, S., Brand, M., and Kawakami, A. (2017). Osteoblast Production by Reserved Progenitor Cells in Zebrafish Bone Regeneration and Maintenance. Dev. Cell. 10.1016/j.devcel.2017.10.015.

8. Dasyani, M., Tan, W.H., Sundaram, S., Imangali, N., Centanin, L., Wittbrodt, J., and Winkler, C. (2019). Lineage tracing of col10a1 cells identifies distinct progenitor populations for osteoblasts and joint cells in the regenerating fin of medaka (Oryzias latipes). Dev. Biol. 10.1016/j.ydbio.2019.07.012.

9. Brown, A.M., Fisher, S., and Iovine, M.K. (2009). Osteoblast maturation occurs in overlapping proximal-distal compartments during fin regeneration in zebrafish. Dev. Dyn. 238, 2922–2928. 10.1002/dvdy.22114.

10. Stewart, S., Gomez, A.W., Armstrong, B.E., Henner, A., and Stankunas, K. (2014). Sequential and opposing activities of Wnt and BMP coordinate zebrafish bone regeneration. Cell Rep. 6, 482–498. 10.1016/j.celrep.2014.01.010.

11. Hou, Y., Lee, H.J., Chen, Y., Ge, J., Osman, F.O.I., McAdow, A.R., Mokalled, M.H., Johnson, S.L., Zhao, G., and Wang, T. (2020). Cellular diversity of the regenerating caudal fin. Sci. Adv. 6, eaba2084. 10.1126/sciadv.aba2084.

12. Jiang, M., Xiao, Y., E, W., Ma, L., Wang, J., Chen, H., Gao, C., Liao, Y., Guo, Q., Peng, J., et al. (2021). Characterization of the Zebrafish Cell Landscape at Single-Cell Resolution. Front. Cell Dev. Biol. 9. 10.3389/fcell.2021.743421.

13. Luecken, M.D., and Theis, F.J. (2019). Current best practices in single-cell RNA-seq analysis: a tutorial. Mol. Syst. Biol. 15, e8746. https://doi.org/10.15252/msb.20188746.

14. Wolf, F.A., Hamey, F.K., Plass, M., Solana, J., Dahlin, J.S., Göttgens, B., Rajewsky, N., Simon, L., and Theis, F.J. (2019). PAGA: graph abstraction reconciles clustering with trajectory inference through a topology preserving map of single cells. Genome Biol. 20, 59. 10.1186/s13059-019-1663-x.

15. Yoshinari, N., Ishida, T., Kudo, A., and Kawakami, A. (2009). Gene expression and functional analysis of zebrafish larval fin fold regeneration. Dev. Biol. 325, 71–81. https://doi.org/10.1016/j.ydbio.2008.09.028.

16. Stelzer, G., Rosen, N., Plaschkes, I., Zimmerman, S., Twik, M., Fishilevich, S., Stein, T.I., Nudel, R., Lieder, I., Mazor, Y., et al. (2016). The GeneCards Suite: From Gene Data Mining to Disease Genome Sequence Analyses. Curr. Protoc. Bioinforma. 54, 1.30.1–1.30.33. 10.1002/cpbi.5.

17. Grotek, B., Wehner, D., and Weidinger, G. (2013). Notch signaling coordinates cellular proliferation with differentiation during zebrafish fin regeneration. Development 140, 1412–1423. 10.1242/dev.087452.

18. Kang, J., Hu, J., Karra, R., Dickson, A.L., Tornini, V.A., Nachtrab, G., Gemberling, M., Goldman, J.A., Black, B.L., and Poss, K.D. (2016). Modulation of tissue repair by regeneration enhancer elements. Nature 532, 201–206. 10.1038/nature17644.

19. König, D., Page, L., Chassot, B., and Jaźwińska, A. (2018). Dynamics of actinotrichia regeneration in the adult zebrafish fin. Dev. Biol. 433, 416–432. https://doi.org/10.1016/j.ydbio.2017.07.024.

20. Stoick-Cooper, C.L., Weidinger, G., Riehle, K.J., Hubbert, C., Major, M.B., Fausto, N., and Moon, R.T. (2007). Distinct Wnt signaling pathways have opposing roles in appendage regeneration. Development 134, 479–489. 10.1242/dev.001123.

21. Smith, A., Zhang, J., Guay, D., Quint, E., Johnson, A., and Akimenko, M.A. (2008). Gene expression analysis on sections of zebrafish regenerating fins reveals limitations in the whole-mount in situ hybridization method. Dev. Dyn. 237, 417–425. https://doi.org/10.1002/dvdy.21417.

22. Akimenko, M.A., Johnson, S.L., Westerfield, M., and Ekker, M. (1995). Differential induction of four msx homeobox genes during fin development and regeneration in zebrafish. Development 121, 347–357.

23. Blum, N., and Begemann, G. (2012). Retinoic acid signaling controls the formation, proliferation and survival of the blastema during adult zebrafish fin regeneration. Development 139, 107–116. 10.1242/dev.065391.

24. Shibata, E., Yokota, Y., Horita, N., Kudo, A., Abe, G., Kawakami, K., and Kawakami, A. (2016). Fgf signalling controls diverse aspects of fin regeneration. Dev. 143, 2920– 2929. 10.1242/dev.140699.

25. Wehner, D., Cizelsky, W., Vasudevaro, M.D., Özhan, G., Haase, C., Kagermeier-Schenk, B., Röder, A., Dorsky, R.I., Moro, E., Argenton, F., et al. (2014). Wnt/β-Catenin Signaling Defines Organizing Centers that Orchestrate Growth and Differentiation of the Regenerating Zebrafish Caudal Fin. Cell Rep. 6, 467–481. 10.1016/j.celrep.2013.12.036.

26. Münch, J., González-Rajal, A., and de la Pompa, J.L. (2013). Notch regulates blastema proliferation and prevents differentiation during adult zebrafish fin regeneration. Development 140, 1402–1411. 10.1242/dev.087346.

27. Li, L., Zhang, J., and Akimenko, M.-A. (2020). Inhibition of mmp13a during zebrafish fin regeneration disrupts fin growth, osteoblasts differentiation, and Laminin organization. Dev. Dyn. 249, 187–198. https://doi.org/10.1002/dvdy.112.

28. Smith, A., Avaron, F., Guay, D., Padhi, B.K., and Akimenko, M.A. (2006). Inhibition of BMP signaling during zebrafish fin regeneration disrupts fin growth and scleroblast differentiation and function. Dev. Biol. 299, 438–454. https://doi.org/10.1016/j.ydbio.2006.08.016.

29. Hoptak-Solga, A.D., Nielsen, S., Jain, I., Thummel, R., Hyde, D.R., and Iovine, M.K. (2008). Connexin43 (GJA1) is required in the population of dividing cells during fin regeneration. Dev. Biol. 317, 541–548. https://doi.org/10.1016/j.ydbio.2008.02.051.

30. McMillan, S.C., Zhang, J., Phan, H.-E., Jeradi, S., Probst, L., Hammerschmidt, M., and Akimenko, M.-A. (2018). A regulatory pathway involving retinoic acid and calcineurin demarcates and maintains joint cells and osteoblasts in regenerating fin. Development 145, dev161158. 10.1242/dev.161158.

31. Ton, Q. V, and Iovine, M.K. (2013). Identification of an evx1-Dependent Joint-Formation Pathway during FIN Regeneration. PLoS One 8, e81240.

32. Geurtzen, K., Knopf, F., Wehner, D., Huitema, L.F.A., Schulte-Merker, S., and Weidinger, G. (2014). Mature osteoblasts dedifferentiate in response to traumatic bone injury in the zebrafish fin and skull. Development 141, 2225–2234. 10.1242/DEV.105817.

33. Hans, S., Kaslin, J., Freudenreich, D., and Brand, M. (2009). Temporally-controlled site-specific recombination in zebrafish. PLoS One 4. 10.1371/journal.pone.0004640.

34. Ornitz, D.M., and Marie, P.J. (2015). Fibroblast growth factor signaling in skeletal development and disease. Genes Dev. 29, 1463–1486.

35. Poss, K.D., Shen, J., Nechiporuk, A., McMahon, G., Thisse, B., Thisse, C., and Keating, M.T. (2000). Roles for Fgf signaling during zebrafish fin regeneration. Dev. Biol. 222, 347–358. 10.1006/dbio.2000.9722.

36. Cudak, N., López-Delgado, A.C., Keil, S., and Knopf, F. (2023). Fibroblast growth factor pathway component expression in the regenerating zebrafish fin. Gene Expr. Patterns, 119307. https://doi.org/10.1016/j.gep.2023.119307.

37. Curado, S., Stainier, D.Y.R., and Anderson, R.M. (2008). Nitroreductase-mediated cell/tissue ablation in zebrafish: a spatially and temporally controlled ablation method with applications in developmental and regeneration studies. Nat. Protoc. 3, 948–954. 10.1038/nprot.2008.58.

38. Singh, S.P., Holdway, J.E., and Poss, K.D. (2012). Regeneration of amputated zebrafish fin rays from de novo osteoblasts. Dev. Cell 22, 879–886. 10.1016/j.devcel.2012.03.006.

39. Bergemann, D., Massoz, L., Bourdouxhe, J., Carril Pardo, C.A., Voz, M.L., Peers, B., and Manfroid, I. (2018). Nifurpirinol: A more potent and reliable substrate compared to metronidazole for nitroreductase-mediated cell ablations. Wound Repair Regen. 10.1111/wrr.12633.

40. Stavri, S., and Zarnescu, O. (2013). The Expression of Alkaline Phosphatase, Osteopontin, Osteocalcin, and Chondroitin Sulfate during Pectoral Fin Regeneration in Carassius auratus gibelio: A Combined Histochemical and Immunohistochemical Study. Microsc. Microanal. 19, 233–242. 10.1017/S1431927612013797.

41. Chen, C.-H., Merriman, A.F., Savage, J., Willer, J., Wahlig, T., Katsanis, N., Yin, V.P., and Poss, K.D. (2015). Transient laminin beta 1a Induction Defines the Wound Epidermis during Zebrafish Fin Regeneration. PLOS Genet. 11, e1005437.

42. Gerber, T., Murawala, P., Knapp, D., Masselink, W., Schuez, M., Hermann, S., Gac-Santel, M., Nowoshilow, S., Kageyama, J., Khattak, S., et al. (2018). Single-cell analysis uncovers convergence of cell identities during axolotl limb regeneration. Science (80-.). 362, eaaq0681. 10.1126/science.aaq0681.

43. Johnson, G.L., Masias, E.J., and Lehoczky, J.A. (2020). Cellular Heterogeneity and Lineage Restriction during Mouse Digit Tip Regeneration at Single-Cell Resolution. Dev. Cell 52, 525–540.e5. 10.1016/j.devcel.2020.01.026.

44. Storer, M.A., Mahmud, N., Karamboulas, K., Borrett, M.J., Yuzwa, S.A., Gont, A., Androschuk, A., Sefton, M. V, Kaplan, D.R., and Miller, F.D. (2020). Acquisition of a Unique Mesenchymal Precursor-like Blastema State Underlies Successful Adult Mammalian Digit Tip Regeneration. Dev. Cell 52, 509–524.e9. 10.1016/j.devcel.2019.12.004.

45. Armstrong, B.E., Henner, A., Stewart, S., and Stankunas, K. (2017). Shh promotes direct interactions between epidermal cells and osteoblast progenitors to shape regenerated zebrafish bone. Development 144, 1165–1176. 10.1242/dev.143792.

46. Poleo, G., Brown, C.W., Laforest, L., and Akimenko, M.-A. (2001). Cell proliferation and movement during early fin regeneration in zebrafish. Dev. Dyn. 221, 380–390. https://doi.org/10.1002/dvdy.1152.

47. Moro, E., Ozhan-Kizil, G., Mongera, A., Beis, D., Wierzbicki, C., Young, R.M., Bournele, D., Domenichini, A., Valdivia, L.E., Lum, L., et al. (2012). In vivo Wnt signaling tracing through a transgenic biosensor fish reveals novel activity domains. Dev. Biol. 366, 327–340. 10.1016/J.YDBIO.2012.03.023.

48. Neumann, C.J., and Nuesslein-Volhard, C. (2000). Patterning of the Zebrafish Retina by a Wave of Sonic Hedgehog Activity. Science (80-.). 289, 2137–2139. 10.1126/science.289.5487.2137.

49. Spoorendonk, K.M., Peterson-Maduro, J., Renn, J., Trowe, T., Kranenbarg, S., Winkler, C., and Schulte-Merker, S. (2008). Retinoic acid and Cyp26b1 are critical regulators of osteogenesis in the axial skeleton. Development 135, 3765–3774. 10.1242/dev.024034.

50. Poss, K.D., Shen, J., and Keating, M.T. (2000). Induction of lef1 during zebrafish fin regeneration. Dev. Dyn. 219, 282–286. 10.1002/1097-0177(2000)9999:9999<::AID-DVDY1045>3.0.CO;2-C.

51. Zheng, G.X.Y., Terry, J.M., Belgrader, P., Ryvkin, P., Bent, Z.W., Wilson, R., Ziraldo, S.B., Wheeler, T.D., McDermott, G.P., Zhu, J., et al. (2017). Massively parallel digital transcriptional profiling of single cells. Nat. Commun. 8, 14049. 10.1038/ncomms14049.

52. Wolf, F.A., Angerer, P., and Theis, F.J. (2018). SCANPY: large-scale single-cell gene expression data analysis. Genome Biol. 19, 15. 10.1186/s13059-017-1382-0.

53. Becht, E., McInnes, L., Healy, J., Dutertre, C.-A., Kwok, I.W.H., Ng, L.G., Ginhoux, F., and Newell, E.W. (2019). Dimensionality reduction for visualizing single-cell data using UMAP. Nat. Biotechnol. 37, 38–44. 10.1038/nbt.4314.

54. Soneson, C., and Robinson, M.D. (2018). Bias, robustness and scalability in single-cell differential expression analysis. Nat. Methods 15, 255–261. 10.1038/nmeth.4612.

55. Lun, A.T.L., McCarthy, D.J., and Marioni, J.C. (2016). A step-by-step workflow for low-level analysis of single-cell RNA-seq data with Bioconductor [version 2; peer review: 3 approved, 2 approved with reservations]. F1000Research 5. 10.12688/f1000research.9501.2.

56. Robinson, M.D., McCarthy, D.J., and Smyth, G.K. (2010). edgeR: a Bioconductor package for differential expression analysis of digital gene expression data. Bioinformatics 26, 139–140. 10.1093/bioinformatics/btp616.

57. Ritchie, M.E., Phipson, B., Wu, D., Hu, Y., Law, C.W., Shi, W., and Smyth, G.K. (2015). limma powers differential expression analyses for RNA-sequencing and microarray studies. Nucleic Acids Res. 43, e47–e47. 10.1093/nar/gkv007.

58. Draper, B.W., Stock, D.W., and Kimmel, C.B. (2003). Zebrafish fgf24 functions with fgf8 to promote posterior mesodermal development. Development 130, 4639–4654. 10.1242/dev.00671.

59. Keil, S., Gupta, M., Brand, M., and Knopf, F. (2021). Heparan sulfate proteoglycan expression in the regenerating zebrafish fin. Dev. Dyn. 250, 1368–1380. 10.1002/dvdy.321.

60. Yoshinari, N., Ando, K., Kudo, A., Kinoshita, M., and Kawakami, A. (2012). Colored medaka and zebrafish: Transgenics with ubiquitous and strong transgene expression driven by the medaka β-actin promoter. Dev. Growth Differ. 54, 818–828. https://doi.org/10.1111/dgd.12013.

61. Deerinck, T.J., Bushong, E.A., Thor, A.K., Ellisman, M., Deerinck, T.J., Bushong, E.A., Ellisman, M., Deerinck, T.J., and Thor, C. (2010). NCMIR methods for 3D EM: a new protocol for preparation of biological specimens for serial block face scanning electron microscopy.

62. Hanker, J.S., Deb, C., Wasserkrug, H.L., and Seligman, A.M. (1966). Staining Tissue for Light and Electron Microscopy by Bridging Metals with Multidentate Ligands. Science (80-.). 152, 1631–1634. 10.1126/science.152.3729.1631.

63. Völkner, M., Wagner, F., Steinheuer, L.M., Carido, M., Kurth, T., Yazbeck, A., Schor, J., Wieneke, S., Ebner, L.J.A., Del Toro Runzer, C., et al. (2022). HBEGF-TNF induce a complex outer retinal pathology with photoreceptor cell extrusion in human organoids. Nat. Commun. 13, 6183. 10.1038/s41467-022-33848-y.

64. Venable, J.H., and Coggeshall, R. (1965). A simplified lead citrate stain for use in electron microscopy. J. Cell Biol. 25, 407–408. 10.1083/jcb.25.2.407.

